# Structure and dynamics of a pentameric KCTD5/Cullin3/Gβγ E3 ubiquitin ligase complex

**DOI:** 10.1101/2023.09.20.558662

**Authors:** Duc Minh Nguyen, Deanna H. Rath, Dominic Devost, Darlaine Pétrin, Robert Rizk, Alan X. Ji, Naveen Narayanan, Darren Yong, Andrew Zhai, Douglas A. Kuntz, Maha U.Q. Mian, Neil C. Pomroy, Alexander F.A. Keszei, Samir Benlekbir, Mohammad T. Mazhab-Jafari, John L. Rubinstein, Terence E. Hébert, Gilbert G. Privé

## Abstract

Heterotrimeric G proteins can be regulated by post-translational modifications, including ubiquitylation. KCTD5, a pentameric substrate receptor protein consisting of an N-terminal BTB domain and a C-terminal domain (CTD), engages CUL3 to form the central scaffold of a cullin-RING E3 ligase complex (CRL3^KCTD5^) that ubiquitylates Gβγ and reduces Gβγ protein levels in cells. The cryo-EM structure of a 5:5:5 KCTD5/CUL3^NTD^/Gβ_1_γ_2_ assembly reveals a highly dynamic complex with rotations of over 60° between the KCTD5^BTB^/CUL3^NTD^ and KCTD5^CTD^/Gβγ moieties of the structure. CRL3^KCTD5^ engages the E3 ligase ARIH1 to ubiquitylate Gβγ in an E3-E3 super-assembly, and extension of the structure to include full-length CUL3 with RBX1 and an ARIH1∼ubiquitin conjugate reveals that some conformational states position the ARIH1∼ubiquitin thioester bond to within 10 Å of lysine-23 of Gβ and likely represent priming complexes. Most previously described CRL/substrate structures have consisted of monovalent complexes and have involved flexible peptide substrates. The structure of the KCTD5/CUL3^NTD^/Gβγ complex shows that the oligomerization of a substrate receptor can generate a polyvalent E3 ligase complex and that the internal dynamics of the substrate receptor can position a structured target for ubiquitylation in a CRL3 complex.

**Significance Statement:** In humans, ∼600 enzyme complexes can carry out protein ubiquitylation, and the most abundant class of these are the cullin3-RING-ligase complexes (CRL3s). CRL3s are multiprotein complexes built around a BTB/cullin3 core, and the incorporation of different BTB proteins into this scaffold results in distinct architectures that ubiquitylate a wide range of substrates. In most cases, it is not known how the complexes are tuned to their substrates. We show that the BTB protein KCTD5 is the central organizer in a CRL3^KCTD5^ complex, and that the architecture and internal dynamics of KCTD5 are essential for positioning a Gβγ substrate protein near an activated ubiquitin for the transfer reaction. This explains how KCTD5 targets Gβγ for proteasomal degradation and regulates cellular activities.

## Introduction

Protein ubiquitylation plays a central role in eukaryotic biology and is often used to regulate intracellular protein levels through proteasomal degradation. The selection of targets for modification is largely determined by the E3 ubiquitin ligases, and the family of cullin-RING E3 ligases (CRLs) catalyze ubiquitin transfer by positioning lysine residues from substrates next to activated E2∼ubiquitin or RING-between-RING (RBR) E3∼ubiquitin conjugates (1, 2). These multi-component modular assemblies use a cullin arm to bridge substrate-binding receptors (SRs) to the RBX1/2 RING component of the ubiquitylation machinery. The CRL3 family, defined by the inclusion of cullin3 (CUL3), can directly engage over 100 different SRs that typically combine a BTB CUL3-binding domain and a substrate-binding domain in a single polypeptide (3). Notably, BTB domains can self-associate into stable dimers, pentamers and oligomers (4–9) and thus drive the multimerization of CRL3 complexes. While multivalency is not a unique feature of the CRL3s (10), most structures of CRL complexes reported to date have involved truncated components that result in monovalent complexes (11, 12).

Proteomic studies have identified KCTD5, KCTD2 and KCTD17, a family of homologous pentameric BTB CRL3 substrate receptors, as interactors of some Gβɣ heterodimers (Gβ_1_ɣ_2_, Gβ_1_ɣ_7_, Gβ_2_ɣ_7_, Gβ_2_ɣ_2_) (13–17), and functional studies have shown that these receptors can promote the ubiquitylation of G protein subunits via CRL3^KCTD^ complexes (17–20). KCTD5 has roles in neurodevelopment and is linked to sleep disorders (21, 22) and members the wider family of 25 human KCTD proteins have been associated with neurological disorders, obesity and cancer through a variety of mechanism (23–26). G proteins, consisting of Gαβɣ heterotrimers and, once activated by GPCRs, the dissociated Gα monomer and Gβɣ heterodimers, activate downstream cellular effectors. Gβγ subunits are involved in a wide range of signalling events in many different subcellular locations (27–29).

CRLs ubiquitylate a wide variety of substrates, ranging from intrinsically disordered proteins anchored via short degron motifs to ordered targets (for example, (30–33)). In the case of ordered substrates such as Gβγ, little is known about how specific lysine residues are selected for modification. Here we show that a CRL3^KCTD5^ complex directly ubiquitylates Gβ_1_ɣ_2_ in a reaction that depends on the RING-between-RING (RBR) E3 ligase ARIH1 (34) and that KCTD5 modulates Gβɣ levels in cells in a proteosome-dependent manner. Cryo-EM structures of KCTD5 in complex with Gβ_1_ɣ_2_ and the N-terminal domain of CUL3 (CUL3^NTD^) show that KCTD5 forms a central pentameric scaffold that simultaneously engages five Gβɣ substrates via its C-terminal domain (KCTD5^CTD^) and five CUL3^NTD^ subunits via its N-terminal BTB domain (KCTD5^BTB^). Internal flexibility is a common feature of E3 ligases, and rotational and translational dynamics between the KCTD5^CTD^/Gβɣ and KCTD5^BTB^/CUL3 moieties of the 5:5:5 complex generate multiple poses of Gβɣ relative to CUL3. We used the experimental structure to model the full reaction complex and show that the internal dynamics of KCTD5 generate conformers in which lysine-23 of Gβ_1_ is positioned near an activated ARIH1∼ubiquitin held by the neddylated CUL3/RBX1 domain, and likely represent the priming complex.

## Results

### KCTD5 assembles a KCTD5/CUL3^NTD^/Gβγ complex

We expressed and purified KCTD5, CUL3^NTD^ and Gβ_1_ɣ_2_, and used tagged KCTD5 to show that CUL3^NTD^ and Gβɣ bind individually and in combination to KCTD5 (Fig. 1*A*). Next, we used bilayer interferometry (BLI) with anchored KCTD5 for more quantitative analyses of the interactions. CUL3^NTD^ binds with a 30 nM dissociation constant, consistent with previous results (6, 8, 35), while Gβɣ binds with micromolar affinity (Fig. 1*B* and SI Appendix, Fig. S1 and Table S1). The presence of CUL3^NTD^ had little effect on the binding affinity of Gβɣ for KCTD5, suggesting little or no cooperativity between the binding of the two partners. Notably, while CUL3^NTD^ and Gβɣ had similar association rates, the dissociation rate for Gβɣ was approximately 60-fold higher than for CUL3^NTD^ and largely accounts for the differences in the affinities between the components. In size exclusion chromatography (SEC) experiments, KCTD5 alone elutes as a single peak consistent with the expected homopentamer seen in crystal structures (36). Additions of CUL3^NTD^ to KCTD5 resulted in the formation of a larger species as detected by SEC, with a maximum shift at a 1:1 molar ratio (SI Appendix Fig. S2). In contrast, the addition of Gβɣ did not produce a complex with KCTD5 that was detectable by SEC, even at a 2-fold molar excess. We attribute this result to the rapid exchange kinetics between KCTD5 and Gβɣ on the time scale of the SEC experiments (∼24 minutes from injection to elution). However, a recent publication showed that KCTD5 and Gβɣ could form a stable complex by SEC (20). Overall,these experiments show that both CUL3 and Gβɣ bind directly and independently to KCTD5, but that Gβɣ interacts more weakly and has a shorter residence time, at least under the tested conditions.

**Fig. 1.**
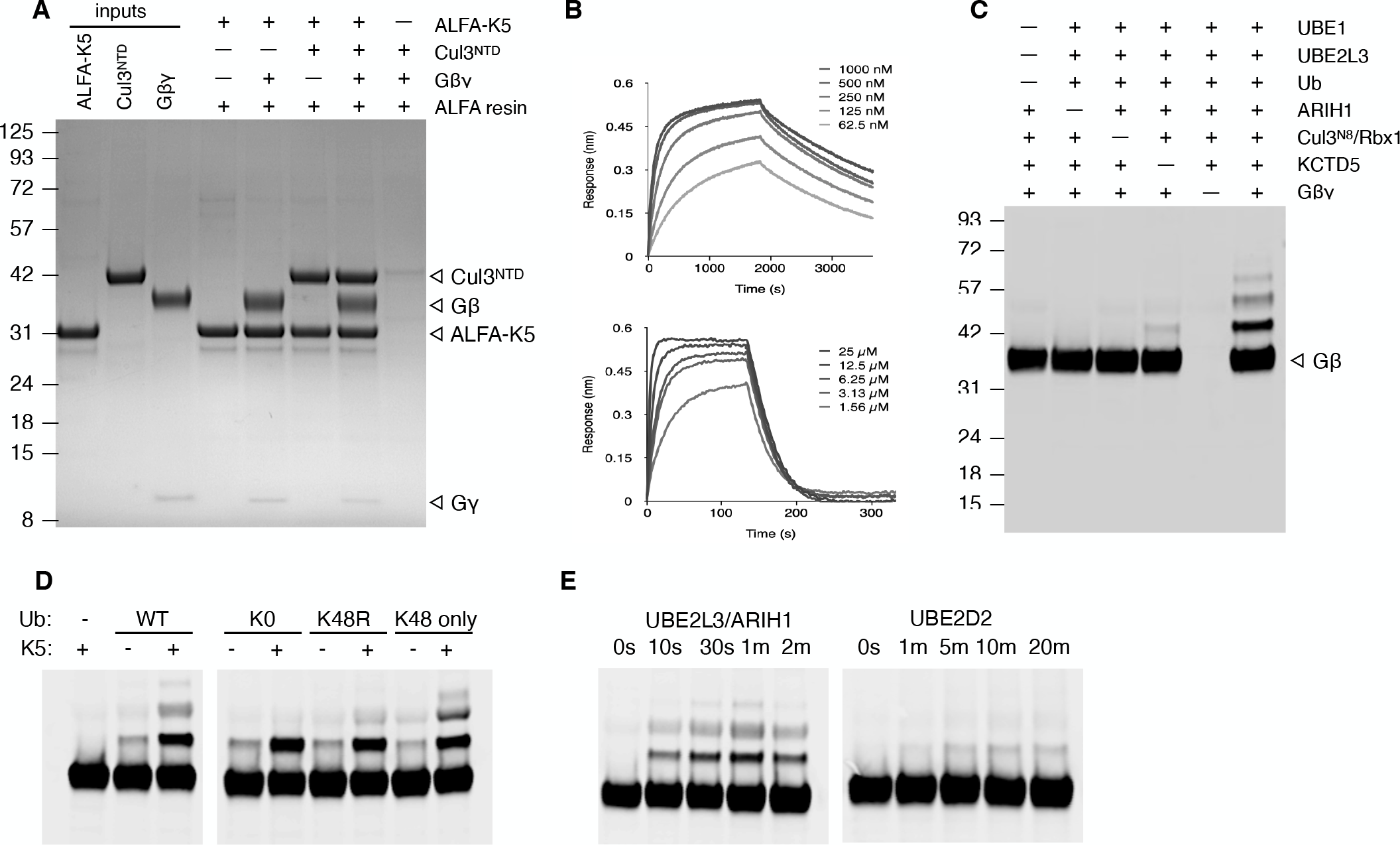
KCTD5 is a CRL3 substrate receptor for Gβγ. (A) Cul3^NTD^ and Gβγ bind directly and independently to ALFA- tagged KCTD5 (ALFA-K5). Coomassie-stained SDS-PAGE. ALFA-resin fractions were eluted with SDS-sample buffer after washing the resin to remove unbound components. (B) Bilayer interferometry (BLI) of sensor-anchored KCTD5 with Cul3^NTD^ (top) and Gβγ (bottom). (C, D, E) anti-Gβ1 Western blots. (C) Cul3^Nedd8^/Rbx1/UBE2L3/ARIH1 ubiquitylates Gβ in a KCTD5 dependent manner (all reactions 2 minutes, 37 °C). (D) CRL3^KCTD5^/ARIH1/UBE2L3 catalyses the ubiqutiylation of a single site on Gβ, and has moderate ubiquitin-K48 chain extension activity. K5: KCTD5; K0: ubiquitin with all seven native lysines replaced with arginines; K48R ubiquitin with a single K48R substitution, K48 only: ubiquitin with six lysine to arginine substitutions, preserving the native lysine at position 48. (E) CRL3^KCTD5^ uses UBE2L3/ARIH1 but not UBE2D2 for the attachment of the initial ubiquitylation of Gβ. All experiments were repeated at least three times and representative data are shown.

### A reconstituted CRL3^KCTD5^ complex ubiquitylates Gβγ

We assembled a defined ubiquitylation system to show that CRL3^KCTD5^ can directly ubiquitylate Gβ subunits, consistent with previous cell-based experiments (17). The reaction requires the E1 protein UBE1, the E2 protein UBE2L3, neddylated CUL3, the RBR E3 ligase ARIH1, and KCTD5 (Fig. 1 *C,D,E*). Thus, CRL3^KCTD5^ uses an E3-E3 super-assembly with ARIH1 for substrate ubiquitylation, as previously shown for several CRL1 complexes (31, 34). The widely used E2 enzyme UBE2D2, which can also directly ubiquitin some substrates (37, 38), was not active under our reaction conditions (Fig. 1E). We were also able to detect the ubiquitylation of Gɣ subunits, but the reaction had a higher background so we focused primarily on Gβ ubiquitylation as a readout for the reaction (SI Appendix Fig. S3). The small amount of Gβ with more than one ubiquitin was due a limited amount of ubiquitin K48 chain formation with UBE2L3/ARIH1, and reactions with a K48R ubiquitin mutant shows that there is a single ubiquitylation site on Gβ (Fig. 1*D*).

### Structure of the KCTD5/CUL3^NTD^/Gβγ complex

KCTD5 has two main folded regions: a BTB domain from residues 45-150 and a CTD domain from 156-209. KCTD5 crystallized under two different conditions revealed that the BTB and CTD domains form independent homopentamers with roughly aligned symmetry axes, and that these can adopt conformational states that differ primarily by a rotation of ∼30° between the N-terminal BTB and C-terminal CTD moieties (36). Molecular dynamics simulations have shown even greater degrees of flexibility between the two domains (39, 20). We determined the structure of a complex containing full-length KCTD5, CUL3^NTD^ and Gβɣ by cryo-electron microscopy. Complexes were prepared *in situ* by directly mixing stock solutions of the three proteins prior to spotting onto cryo-grids and plunge freezing. Initial map construction converged to volumes for either KCTD5^BTB^/CUL3^NTD^ or KCTD5^CTD^/Gβɣ, and maps with density for full complexes could only be obtained by dividing the particles images into subsets, indicative of large-scale internal motions (Fig. 2 and SI Appendix, Figs. S4, S5). Maps generated from the particle image set in which either the bottom KCTD5^BTB^/CUL3^NTD^ region or the top KCTD5^CTD^/Gβɣ region were subtracted (“top only” and “bottom only” maps, respectively) showed improved detail with resolutions of 2.9 Å and 3.5 Å resolution, respectively. Despite minor improvements upon imposing C5 symmetry during map refinements, all final map calculations were carried out without the imposition of symmetry, and we used these subtracted maps to build models for KCTD5^CTD^/Gβɣ and KCTD5^BTB^/CUL3^NTD^ components of the complex. We also obtained maps for multiple conformational states of the entire complex via the subdivided particle image sets (SI Appendix, Fig. S5). A recent report of the structure of a two-component KCTD5/Gβɣ complex required a head-to-tail fusion of KCTD5 with Gɣ to stabilize the assembly for cryo-EM (20). In addition, C5 symmetry was imposed in the refinements. Our structures of the KCTD5/CUL3^NTD^/Gβɣ complex were obtained without artificial cross-links or imposed symmetry and reveal large scale internal motions and pronounced deviations from C5 symmetry. These features have important consequences for the mechanism of substrate ubiquitylation and are described in the sections bellow.

**Fig. 2.**
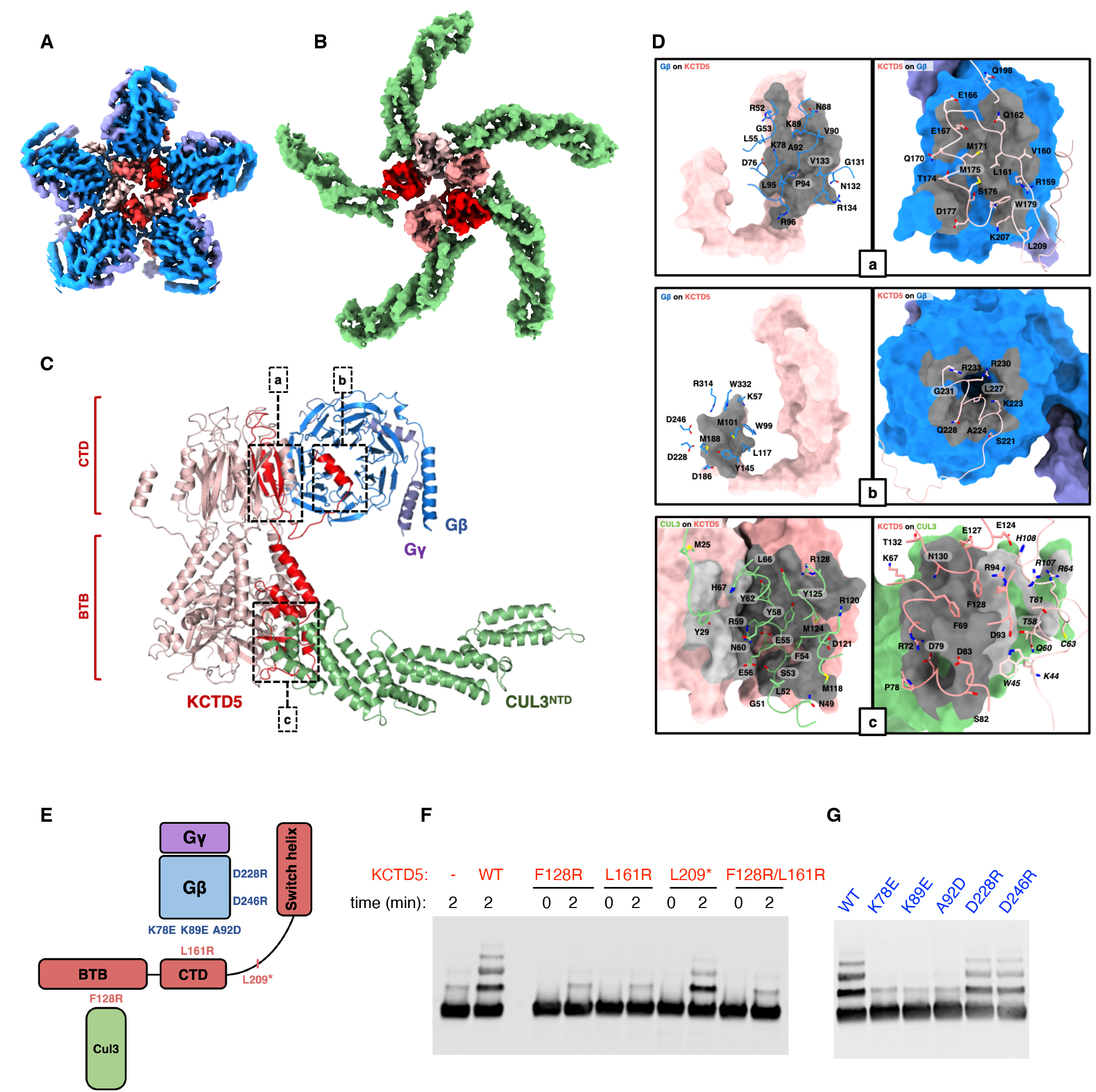
Structure of the pentameric KCTD5/Cul3^NTD^/Gβγ complex. (A) Cryo-em map of the KCTD5^CTD^/Gβγ “top” part of the complex. Densities for KCTD5 are colored different shades of red. Gβ is blue and Gγ is purple. (B) Cryo-em map of the KCTD5^BTB^/Cul3^NTD^ “bottom” part of the complex. Cul3^NTD^ is green. (C) Side view of the model fit to the a map of the intact complex showing only one Gβγ and Cul3^NTD^ chain for clarity. (D) Contact interfaces in the complex as per panel C in an “open book” format with buried surfaces in grey. The primary and distal interfaces from adjacent subunits in “c” are indicated with dark/light shades, respectively. (E) Schematic of the interfaces with key residues indicated. L209* is KCTD5 with a stop codon introduced at residue position 209. (F) Ubiquitylation activity of KCTD5 mutants. (G) Ubiquitylation activity of Gβ mutants. F and G are anti-Gβ Western blots.

### KCTD5^CTD^/Gβɣ subcomplex

In the top half of the complex, each of five Gβɣ units in the pentameric complex interacts with three adjacent chains of the KCTD5 CTD in two contact patches (Fig. 2*A-D*, SI Appendix, Fig. S6 and Movie S1). Five Gβɣ units bind edge-on and project radially from the central KCTD5^CTD^ pentamer. All KCTD5 contacts involve Gβ only and there are no interactions with Gɣ. There are no Gβɣ interactions with KCTD5^BTB^. Remarkably, the two contact patches on Gβ are similar to those engaged by Gα subunits (40) and GRK2 (41) (SI Appendix, Fig. S7) despite the lack of homology between the binding partners. As a result, Gα, GRK2 and KCTD5 binding are likely to be mutually exclusive, suggesting that KCTD5 interacts with free Gβɣ. Remarkably, KCTD12, a pentameric KCTD protein that does not interact with CUL3, uses a C-terminal H-domain to engage 5 Gβɣ units using an unrelated architecture involving different structural elements to regulate GABA_B_ receptors (42). The H1 fold is not compatible the pentamers formed by the KCTD5 CTD. In the major contact between KCTD5 and Gbg (patch “a” in Fig. 2*C,D*), the outermost strand of Gβ blade 1 comprising residues N88-R96 nestles into a shallow groove formed by β1, α1 and β3 from a KCTD5^CTD^ subunit. This surface also includes contacts between L55 and D76 from Gβ blade 1 to residues T169 and S173 in the KCTD5^CTD^ helix from the adjacent subunit. These contacts bury a similar surface on Gβ as the N-terminal helix of Gα in G protein heterotrimers (40). This interaction surface has a small non-polar center (KCTD5 L161, W179 and Gβ A92, P94) flanked by basic residues K78, K89 and R96 on Gβ and acidic residues E167 and D177 on KCTD5.

The second contact patch (“b” in Fig. 3*C,D*) involves residues from the KCTD5 C-terminal tail that follows the BTB CTD, residues 210-234. In the crystal structures of KCTD5, no density was observed for this region, suggesting that these residues in KCTD5 are disordered in the uncomplexed protein (36). The KCTD5 tail is also not included the recent cryo-EM structure of KCTD5 linked to Gβɣ (20). In our structure of the complex, the KCTD5 tail snakes under the long Gβ loop 125-134 and culminates in an α-helix from residues 222-234, which we call the KCTD5 switch helix. The connection from KCTD 214-221 is in weak density, but the Gβ loop 125-134, which is normally flexible in free Gβɣ, makes multiple interchain contacts and is well ordered. Overall, the structure of the KCTD5 C-terminal tail is consistent with the AlphaFold prediction that shows highly exposed residues from 212-221 followed by isolated terminal α-helix (https://alphafold.ebi.ac.uk/entry/Q9NXV2). The basic KCTD5 switch helix is wedged between adjacent Gβɣ units and interacts mainly with the “hot spot” acidic patch of the first Gβ, notably residues D228 and D246, but also forms minor contacts with blade2 of the adjacent Gβ unit. The acidic Gβ hotspot has been observed to bind to a wide range of helical peptides (43), and the KCTD5 switch helix adds to this repertoire.

**Fig. 3.**
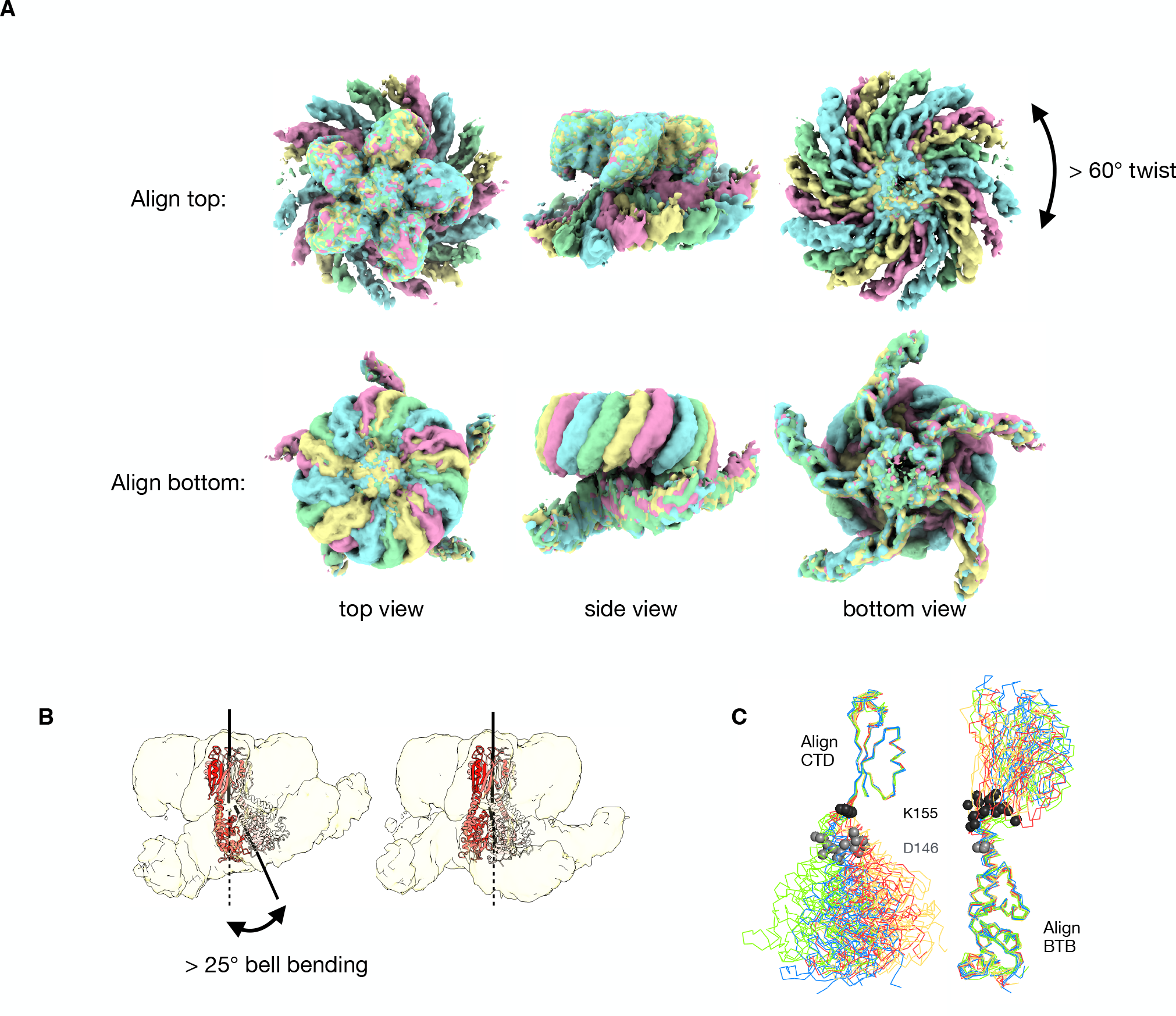
Structural variability in the complex. (A) Alignment of the consensus cryo-em maps for states A, B, C and D (green, cyan, pink, yellow, respectively). Note that the two pseudo-C5 axes between the top and bottom of the complex are not co-linear in these consensus maps. (B) Multi-body refinement reveals additional “bell-bending” variability between the two moieties for each state. (C) Structural alignments between the KCTD5 models built into the consensus cryo-em maps show that the conformational variability in the complex is largely limited to a ∼10 residue region between the BTB and CTD domains. Twenty models corresponding to the five chains of states A, B, C and D are shown, colored according the maps in *A*. The Cɑ atoms of KCTD5 residues 146 and 155 are shown in grey and black spheres, respectively. Bell-binding generates additional variability, but the changes remain localized to the same linker region.

We used structure-guided mutagenesis to test residue substitutions at the KCTD/Gβɣ interfaces (Fig. 2 *E-G* and SI Appendix, Fig. S8, S9 and Table S1). The main contact patch is required for the KCTD5/Gβɣ association, as mutations L161R in KCTD5 and K78E, K89E or A92D in Gβ disrupt binding without affecting the fold of either protein or CUL3^NTD^ association. Accordingly, these mutations also fully disrupt the ubiquitylation activity of the complex. In contrast, deletion of the KCTD5 tail via mutation L209* reduces the Gβɣ binding affinity by approximately 10-fold. Basic substitutions in Gβ at positions D228 or D246 or KCTD5 mutant L209* reduce but do not fully disrupt activity, demonstrating that the KCTD5 tail is of secondary importance in stabilizing the complex.

Consensus refinement of CTD/Gβɣ with no imposed symmetry restrains reveals deviations from C5 symmetry in this part of the complex. The angles between the adjacent Gβɣ subunits range between 70° and 74° with an average of 72° as expected for 5-fold symmetry.

Notably, the sandwiched KCTD5 C-terminal helix density is weaker when the Gβɣ arms are closer together and stronger when they are farther apart, suggesting that a fully engaged KCTD5 helix pushes the Gβɣ units apart from the ideal 72° spacing. In other words, it appears that the pentameric complex cannot accommodate 5 fully engaged terminal helices at the same time, indicating a degree of frustration in the complex. A 3DFlex analysis (44) of the dynamics in the complex shows that the outer edges of the Gβɣ units, which include the N terminal helices from Gβ and Gɣ, are more dynamic than the central core near the pseudo-C5 axis (Movie 1).

### KCTD5^BTB^/Cul3 subcomplex

In the bottom part of the complex, the CUL3^NTD^ interacts with two adjacent KCTD5^BTB^ domains (contact patch “c” in Fig. 2 *C,D*), consistent with earlier modeling studies (6, 8) and a recently determined KCTD7 structure (20). All reported BTB/CUL3, SKP1/CUL1, ELOC/CUL2 and ELOC/CUL5 structures involve a common primary interface and a second smaller distal interface that varies widely between substrate adaptor/cullin pairs (45, 4, 6). A major component of the primary interface is the α2/v-loop/β3 region from the common BTB/SKP1/ELOC fold (6, 45) (SI Appendix, Fig. S10), which consists of residues F69-R94 in KCTD5. These elements contact H2, H4 and H5 in the cullin-repeat 1 (CR1) domain of their cognate cullins and include residues that are displaced by CAND1 to promote SR exchange (46, 47). In the dimeric KLHL and SPOP BTB families, the v-loop includes the critical φ-X-E motif (4, 48), but this motif is not found in the v-loops of the KCTD family (6), even though members of both families bind to CUL3 with nanomolar affinity. In the adjacent distal surface, the short KCTD5 helix α1 (R59- R64) on the adjacent BTB subunit fits into a shallow groove between CUL3 helices H1 and H2. In an apparent example of convergent evolution, the distal interface engages non-homologous elements from the 3-box region of the BACK domain from the same chain in KLHL/SPOP, or from the F-box or BC-Box in SKP1 and ELOC, respectively (SI Appendix, Fig. S10). As expected, mutation at KCTD5 position F128 in the BTB domain disrupted the interaction with CUL3^NTD^ (6) but not with Gβɣ, and abolished the ubiquitylation activity of the complex (Fig. 2 *E, F* and SI Appendix, Fig. S8, S9 and Table S1).

Pentameric KCTD^BTB^ rings often deviate from C5 symmetry, and “ring opening” has been observed in KCTD BTB domains (6–8). This is recapitulated in the bottom-only consensus cryo-EM map in which there are no contacts between one pair of adjacent BTB domains (Fig. 2B). A 3DFlex (44) analysis of the KCTD5^BTB^/CUL3^NTD^ subcomplex reveals structural states ranging from a closed BTB pentamer to an open ring (Movie 1). One consequence of these dynamics is that the CUL3^NTD^ chain at the open ring position engages KCTD5 via the primary interface only and does not make the distal contact (SI Appendix, Fig. S11). Notably, the map density for this CUL3 chain is weaker than for the other four chains, suggesting partial occupancy for this subunit due to reduced affinity.

### Structure and dynamics in the intact KCTD5^BTB^/Cul3/Gβɣ complex

We were able to obtain maps for the entire complex only by subdividing the cleaned particle image set into subgroups; these maps differed in the relative positions of the top and bottom of the complex. Larger number of subgroups (with smaller numbers of particle images in each) lead to finer differences between the maps, suggesting continuous motions in the complex rather than discrete heterogeneity. We tested different methods to analyze heterogeneity in this dataset (44, 49, 50) with full and subdivided particle image sets. We found that multi-body refinement (49) on four subsets was best suited to the current problem and yielded clear improvements for both moieties of the structures. We first generated four ∼4 Å maps from consensus refinements on particle image subsets A, B, C, D; these had unambiguous density for the intact complexes but still had significant smearing in the KCTD5^BTB^/CUL3^NTD^ regions, and we combined the top and bottom focused maps to improve these densities (Table S2). For each of the four substates, we generated 10 composite multi-body maps for a total of 40 maps (Fig. 3 and SI Appendix, Fig. S5). In principle, map density at the interfaces between bodies is ill-defined, but we were able to unambiguously determine the connectivity between the BTB and CTD domains despite weaker density in these regions (SI Appendix, Fig. S12).

The resulting models differ mainly by on-axis rotations of over 60° (“twist”) between the two pseudo-C5 moieties and by rotations of up to 25° about an axis perpendicular to the pseudo-C5 axes (“bell-bending”) (Movie 2). The consensus maps A, B, C and D mostly capture the twisting component of the ensemble of states, while the multi-body component maps mostly reveal bell-bending in each state. There are also on-axis translations of up to 5 Å. There are few contacts between the two halves of the complex, and CUL3^NTD^ and Gβɣ come into direct contact only at higher bell-bending angles.

Essentially all of the structural variability in the complex can be localized to a 10-residue connection between the BTB and CTD domains of KCTD5 (Fig. 3*C*), and this is recapitulated in an analysis of the crystal structures of uncomplexed KCTD5 (36) (SI Appendix, Fig. S13). The hinge region begins at residue D146 in the last α-helix of the BTB domain and ends at K155 prior to the first β-strand of the CTD. The flexibility in the hinge, which accounts for twist, bell- bending, and translation, appears to be evenly distributed over the 10 residues. KCTD5 homologs have low sequence conservation in this region, apart from a composition of mostly hydrophilic amino acids (SI Appendix, Fig. S13).

### CRL3^KCTD5^ positions Gβγ for ubiquitylation

We generated models of intact CRL3^KCTD5^ complexes by extending CUL3 to include structures of the ubiquitylation machinery assembled at the C-terminal half of the cullin. Modeling was based on the CUL1/RBX1/ARIH1(Rcat)∼ubiquitin components of the TS2 state of a CRL1^FBXW7^ structure (31). In the model, all five CUL3 arms arc towards the CTD of the KCTD5 pentamer in a suprafacial arrangement (11) while allowing just enough space for the inclusion of Gβɣ. The arms are positioned such that all five ubiquitin C-termini face the central pseudo-C5 axis of the pentamer and do not produce unfavorable clashes over the ensemble of conformations (Fig. 4 and Movie 2). The twist and bell-bending motions between the two moieties of the complex generate a “cloud” of ARIH1(Rcat)∼ubiquitin reactive sites that populate a solid toroid that surrounds the Gβɣ surface (Fig. 5B). Some of the structural states result in close approaches between the activated ubiquitin C-termini and the Gβɣ units, notably surface exposed lysines K23 on Gβ and K29, K32 and K46 on Gɣ (Fig. 5B and SI Appendix, Fig. S14). Of these, Gβ K23 is the most conserved across the human isoforms (SI Appendix, Fig. S15). Although only a fraction of the conformational states are poised for ubiquitin transfer to Gβ K23, this appears to be sufficient for the reaction, as CRL3^KCTD5^ ubiquitylates Gβ exclusively at K23 (Fig. 4C and SI Appendix, Fig. S16). It is important to note that the ARIH1(Rcat)∼Ub transferase module is highly dynamic relative to the cullin N terminal domain (31), and these additional degrees of freedom would further expand the volume of the reactive cloud.

**Fig. 4.**
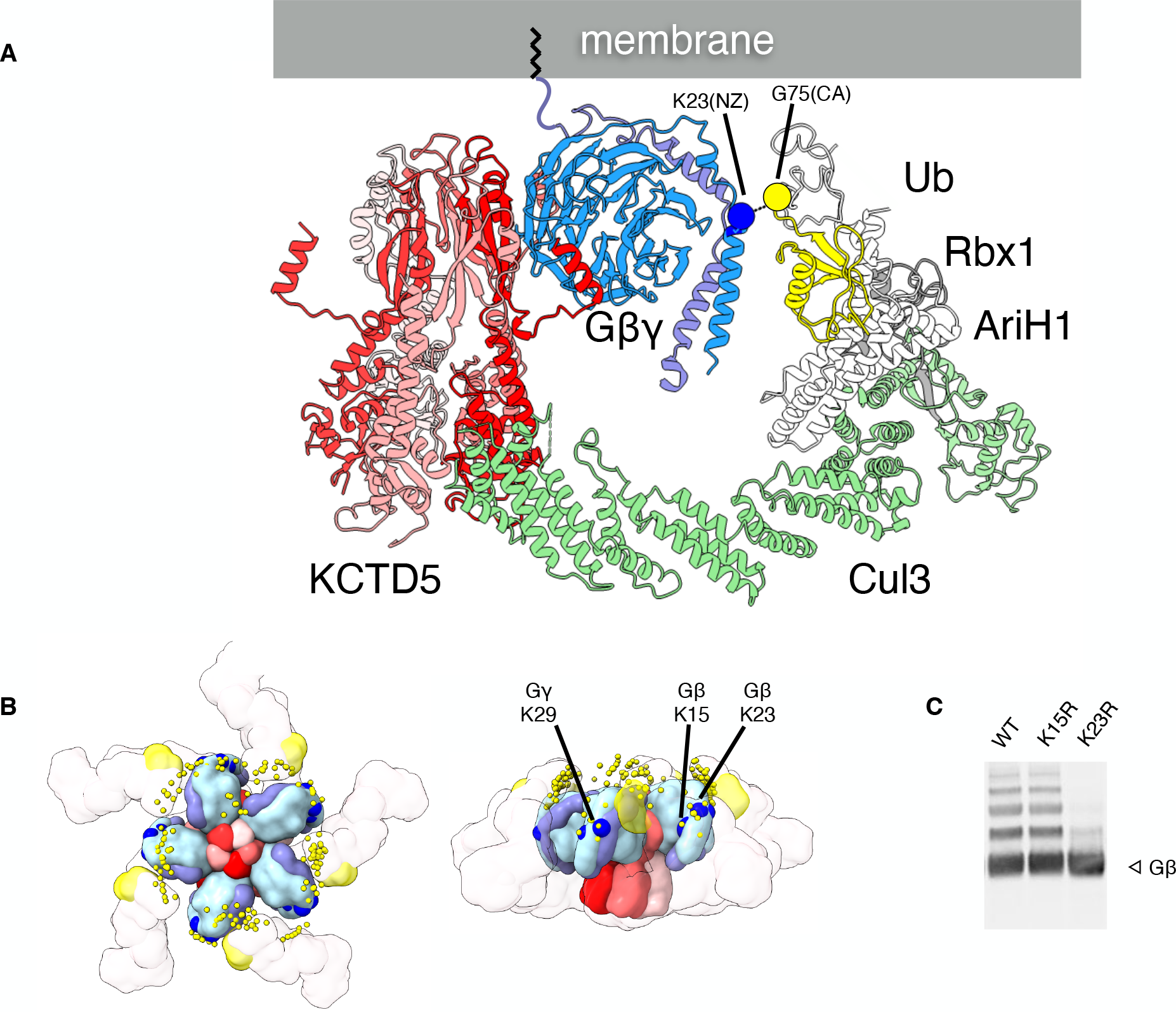
Model of an active CRL3^KCTD5^ complex. (A) Side view of conformational state B of the KCTD5/Cul3^NTD^/Gβγ complex with one CUL3 arm extended based on the TS2 state of the CRL1^FBXW7^ complex (PDB ID 7B5M). Rbx1 is in grey, AriH1 (Ariadne and Rcat domains) is in white, and ubiquitin is in yellow. The conformational state shown here places the NZ atom of Gβ K23 within 9 Å of the ubiquitin G75 CA atom with an unobstructed path to the thioester bond of the ubiquitin/AriH1∼ubiquitin conjugate. The structure of the complex is consistent with membrane-anchored Gβγ. (B) Surface representation of one conformer of the extended complex. The small yellow circles indicate the positions of the ubiquitin G75 Cα in the ensemble of the conformations of the complex, based on MultiBody maps for states A-D. C. Ubiquitylation of Gβ WT and mutants (anti-Gβ Western blot).

**Fig. 5.**
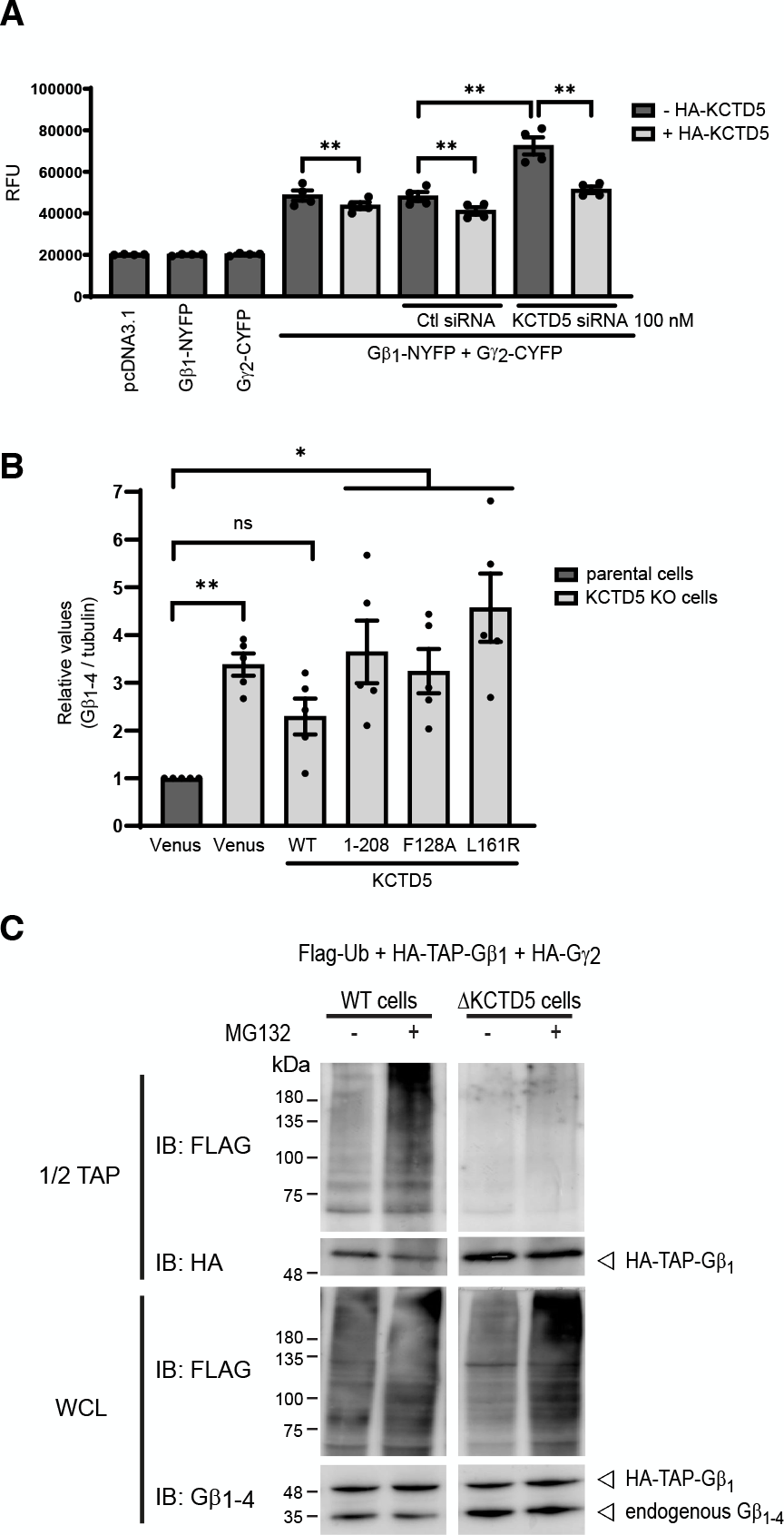
Reducing KCTD5 levels alters Gβγ expression and ubiquitination patterns. (A) The level of Gβγ is altered with KCTD5 knockdown or overexpression. BiFC was measured between the split YFP fusion proteins Gb1-NYFP_(1-158)_ and Gg2-CYFP_(159-238)_, in presence or absence of overexpressed HA-KCTD5 or 100 nM siRNA. Fluorescence intensities are presented as mean +/- SEM for 4 different experiments. Data were analysed by two-way ANOVA followed by Tukey’s multiple comparisons test where ** p < 0.01. (B) Pan-Gβ_1-4_ expression measured by Western blot in parental and KCTD5 KO cell lines stably expressing either the WT, the C-terminal truncated (1-208*), F128A or L161R KCTD5 mutants. For the sake of comparison, both parental and KCTD5 KO cell lines expressed a non-KCTD5 control protein (Venus, a YFP). Bars represent the average values from 5 independent experiments expressed as relative values normalized to expression of β-tubulin. Error bars are SEM. Statistical analysis was done by one-way ANOVA followed by a Dunnett post-hoc test, * p<0.05, **p<0.01 and ns means non-significant. (C) KCTD5 levels alter ubquitylation of Gβγ and their interacting proteins. Western blot analysis from parental and KCTD5 KO lines expressing Flag-Ubiquitin (Ub), HA-TAP-Gβ1 and HA-Gγ2 that were treated or not with 10 mM MG-132 for 8 hours, lysed and subjected to streptavidin purification. Blots are representative of 3 independent experiments.

### KCTD5 regulates Gβγ levels in cells

Validating the KCTD5/Gβɣ interaction from (14), we showed that KCTD5 could be pulled down with TAP-Gβ1 and not with TAP-Smad2 (SI Appendix, Figure S17). Pulling down KTCD5- FLAG, we also show that the co-immunoprecipitation was bidirectional. We next used bimolecular fluorescence complementation between Gβ and Gɣ subunits and examined the effects of knocking down KCTD5 in HEK 293 cells. Knocking down KCTD5 with siRNA increased the levels of Gβɣ pairs (Fig. 5A) and were reduced by overexpressing siRNA-resistant KCTD5. Next, to confirm effects on native Gβ levels, we used Western blotting with an antibody the recognizes Gβ1-4 in parental cells and in a line of HEK 293 cells where we knocked out KCTD5 using CRISPR/Cas9. In KCTD5 KO cells, the levels of Gβ are significantly increased compared to parental cells (see column 1 versus 2 in Fig. 5B). Restoring expression of KCTD5 WT lowered the levels of detected Gβ, but KCTD5 mutants L209*, F128A and L161R did not (Figure 6B). These mutants disrupt Gβɣ binding, although the deletion of the KCTD5 C-terminal tail in L209* retained some biochemical activity (Fig. 2 and SI Appendix, Fig. S1, S8, S9 and Table S1). Next, we examined the ubiquitylation of Gβ using TAP tagging. Gβ was pulled down in parental cells, with and without treatment with the proteasome inhibitor MG-132 (Fig. 5C, left panels). MG-132 treatment increases Gβ ubiquitylation, demonstrating the involvement of the 28S proteasome in regulating Gβ levels. In contrast, very little ubiquitylation is seen in the KCTD5 KO cells and levels of Gβ are increased but insensitive to MG-132 treatment (Fig. 5C, right panels), concordant with the role for KCTD5 as a substrate receptor for Gβ regulation by CUL3-mediated proteasomal degradation.

**Fig. 6.**
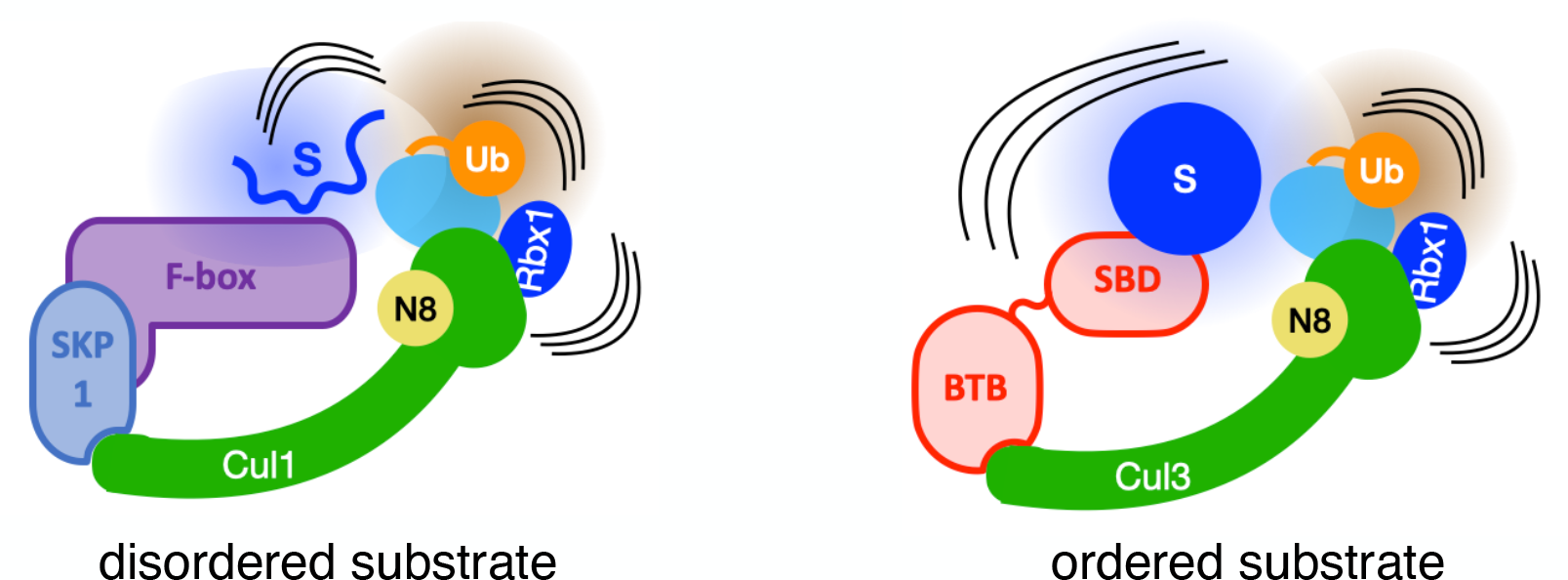
Model for the role of CRL dynamics in positioning an activated ubiquitin C-terminus near to a target substrate. See text for details.

## Discussion

The CRL framework depends on the internal dynamics between multiple structural blocks to adapt to a variety of substrates. For example, there are large reorganizations of the UBE2D∼ubiquitin or ARIH1(Rcat)∼ubiquitin transferase modules relative to the substrate scaffolding module over the catalytic cycle (30, 31). At the substrate end, intrinsically disordered degrons or phosphodegrons can occupy an extended, fuzzy volume (51), and ubiquitylation can occur only when the volumes sampled by the activated ubiquitin thioester and substrate nucleophiles intersect (52). This is a necessary condition, but additional factors, such as nucleophile positioning, are important. With ordered substrates, the requirements for close approach and proper orientation are likely more difficult to achieve and can be expected to be case-specific. The CRL3^KCTD5^ system described here includes large scale internal motions in the KCTD5 substrate receptor within a pseudo-C5 pentamer. The partial decoupling of the KCTD5^CTD^/Gβɣ block from the KCTD5^BTB^/CUL3^NTD^ block allows for the close approach of the rigid Gβɣ substrate to the ubiquitin transferase module (Fig. 6), and we propose that this flexibility is important for catalytic activity. The architecture of the complex is remarkably tuned to the substrate – the shape and position of Gβɣ on KCTD5^CTD^, the geometry of the KCTD5^BTB^/CUL3 interface, the arc of the cullin, and the position of the ARIH1∼Ub on the modelled transferase module – all contribute to the favorable placement of the active groups with little evidence of steric clashes.

The CRL3^KCTD^ reaction complex we describe here is compatible with membrane-bound Gβγ (Fig. 4), leading to a model where the recruitment of pentameric KCTD5 nucleates a cluster of Gβγ and CUL3 complexes at inner membrane surfaces. KCTD5-driven ubiquitylation can modulate Gβγ levels in cells (Fig. 5 and (18–20)), and it will be important to determine how this activity is regulated. Possible mechanisms include post-translational modifications (although none are apparently required for substrate binding), component concentrations, and subcellular localizations. The existence of sub-stoichiometric CRL3^KCTD5^ complexes with incomplete occupancy would lead to unproductive states, and activity may depend on threshold effects due to the concentrations and availability of limiting factors. The fast substrate exchange kinetics and slow CUL3 dissociation rates we describe here have been observed in other CRL complexes (53). Component availability and exchange rates, both spontaneous and catalyzed, are widely used strategies in the regulation of CRL1 networks (46), and we expect these effects to be at work in CRL3^KCTD^ networks as well.

The KCTD family of proteins are involved in a wide range of biological processes and participate in a variety of cellular signaling pathways (23–25). Members of clades E (KCTD2/5/17) and F (KCTD8/12/16) use distinct binding modes to engage Gβγ subunits and downregulate GPCR signaling though CUL3-dependent and CUL3-independent mechanisms, respectively (SI Appendix, Fig. S7 and (16–19)). Published studies of phenotypic effects of KCTD proteins demonstrate a remarkable plasticity which likely results from different levels or subtypes of KCTD proteins present in the different cell models used. The regulation of the levels or activity of distinct populations of Gβγ subunits via KCTD proteins provides an rationale for the pleiotropic effects in distinct cells, as Gβγ subunits are central players in the organization and regulation of cellular signaling (27, 28, 54).

## Materials and Methods

### Protein expression and purification

Expression plasmids and mutants for KCTD5, Gβ_1_, Gɣ_2_, CUL3, and RBX1 were made by gene synthesis (Genscript) unless noted otherwise. Human KCTD5 was expressed as thioredoxin-6His-(TEV)-AviALFA-(3C)-KCTD5 or 6xHis-SUMO- (TEV)-AviALFA-(3C)-KCTD5 from pET32 vectors in *E. coli* (the bracketed terms indicate cleavage sites for TEV and 3C proteases, respectively). Cul3^NTD^ (residues 1-381), UBA1, UBE2L3 and ARIH1^91-557^ were expressed as N-terminal His-tagged fusions from pMCSG7, pET28, pET28aLIC, and pProExHTb vectors in *E. coli*, respectively. *E. coli* expressed proteins were purified by Ni-NTA affinity chromatography and tags were removed by TEV, 3C or thrombin digestion. Proteins were further purified exchanged by size exclusion chromatography into buffer A (25 mM HEPES pH 7.5, 100 mM NaCl, 0.1 mM TCEP, 5% glycerol) prior to concentration and flash freezing in liquid nitrogen.

Gβ_1_ɣ_2_ and CUL3/RBX1 were prepared by co-expression in HEK 293 cells. Gβ_1_ was cloned into a piggyBac vector (PB) (55) without any tags. Full-length Gγ_2_(C68S), a prenylation-deficient variant, was inserted into a PB vector downstream of GFP(3C). Plasmids for Gβ_1_ and GFP-Gγ_2_ were mixed with PBase and PB-RB vectors (55) and transfected into HEK 293 GnT1^-/-^ cells. Stable cell lines were selected for resistance to puromycin and blasticidin and scaled up in FreeStyle suspension cultures. Expression was induced with doxycycline and cells were harvested after 48- 72 hours. GFP-tagged proteins were purified from cell lysates on GFP affinity columns and eluted with 3C protease. Proteins were further purified exchanged by size exclusion chromatography into 25 mM HEPES pH 7.5, 100 mM NaCl, 0.1 mM TCEP and 5% glycerol prior to concentration and flash freezing in liquid nitrogen. All Gβɣ proteins in this study were in the Gβ_1_ɣ_2_(C68S) background. Proteins were further purified exchanged by size exclusion chromatography into buffer A prior to concentration and flash freezing. GFP(3C)-CUL3 and untagged RBX1 were co-expressed in HEK 293 cells and purified as described for Gβɣ, except that the protein was neddylated on-resin with NAE1/UBA3, UBE2M, and NEDD8 (R&D Systems) prior to 3C elution.

### Size exclusion chromatography

Analytical size exclusion chromatography was carried out on a Waters HPLC with an ENrich^TM^ SEC 650 10 x 300 column (Bio-Rad) in 20 mM HEPES pH 8.0, 150 mM NaCl, 0.1 mM TCEP at a flow rate of 1.0 mL/min.

### Co-purification binding assay

ALFA-tagged KCTD5 was added to anti-ALFA nanobody resin in buffer A and washed three times. Cul3^NTD^ and/or Gβγ were added and incubated at 4 °C for 15 min, followed by a second wash and elution into SDS sample buffer. Samples were analyzed by SDS/PAGE and visualized with Coomassie staining.

### *In vitro* ubiquitylation assay

We adapted a pulse-chase ubiquitylation assay (34) for CRL3 complexes. Ubiquitin-charged UBE2L3 or UBE2D2 (E2∼Ub) were generated with 0.3 µM mUBA1, 10 µM UBE2L3 or UBE2D2, and 10 µM ubiquitin in 25 mM HEPES pH 7.5, 150 mM NaCl, 5 mM MgCl_2_ and 1 mM ATP at 37 °C for 15 min and quenched with 10 U/µL apyrase. Reactions were initiated by mixing equal volumes of E2∼Ub with neddylated CUL3/RBX1, KCTD5, Gβ_1_γ_2_, and ARIH1 for UBE2L3∼Ub. Reactions were stopped with SDS-PAGE sample buffer and analyzed by Western blotting with anti-Gγ_2_ (7-RE20, Santa Cruz sc-134344) and anti- Gβ_1_ (Sigma-Aldrich sab2701168).

### Biolayer Interferometry

An Octet RED384 system (Sartorius) instrument was used for biolayer interferometry. Thioredoxin-fusion KCTD5 was anchored onto anti-Penta-His (HIS1K) biosensors and dipped into wells containing CUL3^NTD^ or Gβγ in 20 mM HEPES pH 8, 150 mM NaCl, 1 mM TCEP, 0.05% Tween-20. A different sensor was used for each CUL3^NTD^ concentration because of the slow off-rate for this analyte. Instrument software was used to process and fit the data to a 1:1 kinetic model and a steady-state model.

### Cryo-electron microscopy and model building

Cryo-EM samples were prepared by incubating a mixture of 12 µM KCTD5, 18 µM Cul3^NTD^, and 18 µM Gβγ in 25 mM HEPES pH 8.0, 150 mM NaCl, 0.1 mM TCEP at 4 °C for 30 min. The solution was applied to glow-discharged homemade nanofabricated holey gold grids (56) in a Vitrobot Mark IV and vitrified by plunging into liquid ethane. High-resolution data were collected on a Titan Krios G3 electron microscope (Thermo Fisher Scientific) operated at 300 kV equipped with a prototype Falcon 4 direct detector device camera. Movies were collected at a nominal magnification of 75,000×, corresponding to a calibrated pixel size of 1.03 Å, and a total exposure of ∼49.3 e^-^/Å^2^ over 40 frames. Data from untilted movies and 40° tilted movies were processed with cryoSPARC (57). Masks of the initial “bottom” and “top” moieties were used in particle subtraction to generate top-only and bottom-only particle image datasets. Ab initio maps were calculated from the subtracted particle sets and used for nonuniform refinement (58). Final maps were sharpened with DeepEMhancer (59). Models for KCTD5 (3DRX), CUL3^NTD^ (4EOZ) and Gβγ (1GP2), were docked into the maps and rebuilt and refined with COOT (60) and Phenix (61) to give final coordinates for KCTD5^CTD^/Gβγ and KCTD5^BTB^/CUL3^NTD^.

The unsubtracted particle image set was subdivided into sets A, B, C and D with cryoSPARC heterogenous refinement and imported to Relion (62) for de novo map generation and initial 3D refinement. We improved these maps in two ways. In the first method, we used Relion multi-body refinement (49, 63) with the four Relion consensus maps and masks for the top and bottom bodies. In each of the four cases, the second most abundant component reflected bell-bending and was selected for further analysis. Ten multi-body composite maps for each state were used to build a total of 40 models with Namdinator (64) and further refined with Phenix. The forty multi-body maps and models have been deposited at Zenodo. In the second method, we used phenix.combine_focused_maps with the four Relion consensus maps as templates and the top- only and bottom-only focused maps. The four maps and models from this method have been deposited to the EMDB and PDB (Table S2). Figures were prepared with ChimeraX (65) and Pymol (66). Software used in this project was curated by SBgrid (67).

### Cell culture and transfection

HEK 293 cells were grown in DMEM high glucose, supplemented with 5% fetal bovine serum and penicillin/streptomycin (100 U/ml and 100 μg/ml, respectively, Wisent), at 37°C under humidified 5% CO_2_ environment. HEK 293 cells were transfected using Lipofectamine 2000 reagent (Invitrogen). Five hours after addition of DNA:transfection reagent mixes, media was replaced for complete cell culture media and maintained for 48 hours before performing experiments. Constructs were verified by Sanger sequencing (Genome Québec).

### Bimolecular Fluorescence Complementation (BiFC)

HEK 293 cells in 6-well plates were transfected with plasmids coding for split YFP-tagged fusion proteins expression vectors Gβ1- NYFP_(1-158)_ and Gγ2-CYFP_(159-238)_, with pMAX(-)HA-KCTD5 (kind gift from Dr Goldstein) or pcDNA3.1(+) as a negative control, along with 100 nM non-targeting control siRNA or 100 nM human KCTD5 siRNA (Dharmacon, (68)). Forty-eight hours after transfection, cells were washed with 1x PBS containing 0.1 % glucose, detached and transferred to 1.5 ml tubes, and spun down for 2 minutes at 400 g. Cell pellets were resuspended in 100 μl of 1x PBS containing 0.1 % glucose, and transferred to a white 96-well microplates (white Optiplate; Perkin Elmer Life and Analytical Sciences). Fluorescence was assessed using a BioTek Synergy 2 multi-mode plate reader using a 485/20 nm excitation filter and a 528/20 nm emission filter. Statistical analyses were performed using GraphPad Prism9.

### Generation of KCTD5 knockout cell line using CRISPR/Cas9 mediated genome editing

A guide RNA targeting the human KCTD5 gene close to the methionine start site was designed with the lowest off-target activity. The sgRNA was generated by annealing two complementary oligos: sgRNA-hKCTD5 sense CACCGCGAGCTCCTGTCGCCGGCC and sgRNA-hKCTD5 anti-sense AAACGGCCGGCGACAGGAGCTCGC followed by phosphorylation of the double-stranded DNA by T4 kinase (NEB). The sgRNA was cloned into pX330 (a generous gift from S. Angers, University of Toronto) at the BbsI restriction site. Insertion was confirmed by Sanger sequencing.

The day before transfection, HEK 293 cells were plated in 6-well plates and transfected as described above. Forty eight hours later, cells were detached by trypsinization (Wisent), counted using a TC20 cell counter (Bio-RAD) and diluted in complete media to 5 cells/ml. 100 μl of diluted cells were aliquoted into 96-well plates to 0.5 cell/well and left to expand for 2 weeks until confluent. Cells were collected and expanded in T75 flasks to confluency. Aliquots of the cells were frozen and remaining cells were pelleted by centrifugation at 500g for 5 minutes and genomic DNA was extracted and purified (GeneAid).

Screening of clones for both edited alleles was done by RFLP, as the gRNA overlapped a NaeI restriction site. The genomic sequence surrounding the Cas9 edited site was amplified by PCR (Phusion High-Fidelity DNA polymerase, Thermo) including both forward: gaaggctagggtcgaggtctg and reverse primers: catccttgtctgagtccaggtc. PCR reactions were purified using GeneAid and ∼100 ng of DNA subjected to digestion by NaeI (NEB). Clones that showed no apparent digestion product (due to the modification of the NaeI restriction site) were subjected to amplicon sequencing and clone D2 showed that both alleles were mutated creating a frameshift: one allele with an extra C and the other bearing a 13 nucleotide deletion.

### Rescue experiments

The pcDNA3.1(-)zeo-full-length KCTD5 (1-234) and the C-terminal truncated mutant (1-209*) were generated by PCR using a common forward primer XhoI -HA- hKCTD5: gatcctcgaggccaccatgtacccgtacgac and reverse primers HindIII-full length HA-hKCTD5 (1-234): GATCAAGCTTTCACATCCTTGAGCCTCGTTCTTGC and HindIII-truncated HA- hKCTD5 (1-208): GATCAAGCTTTCACTCCTTGGACACCACACAGAGGAAC. PCR was performed using the template pMAX(-)-HA-KCTD5. Inserts were digested with XhoI and HindIII (NEB) for 1 hour at 37 °C and ligated into pcDNA3.1(-) zeo (Invitrogen) using the same restriction sites. F128A and L161R mutants were made using PCR-based site-directed mutagenesis using the forward: GTTGGAGGAAGCAGAAGCTTACAATATCACCTC and reverse oligonucleotides: GAGGTGATATTGTAAGCTTCTGCTTCCTCCAAC for F128A and forward:

CATGTGTACCGTGTGCGGCAGTGCCAGGAGGAG and reverse oligonucleotides: CTCCTCCTGGCACTGCCGCACACGGTACACATG for L161R.

Parental and KCTD5 KO lines were cultured and transfected with wt or mutant HA-tagged KCTD5; some parental and KO cells were transfected with yellow fluorescent protein (Venus). The following day, cells were replated in 10 cm dishes and left to grow until the next day. 100 μg/ml of zeocin (Invivogen) was used to select cells stably expressing proteins of interest. Lines were expanded, frozen and verified by western blot (**data not shown**).

### Immunodetection analysis

Cells grown in 6-well plates were washed once with 2 ml of PBS and directly frozen at -80 °C. Total cell lysates were prepared by collecting cells in 250 μl of ice-cold lysis buffer (50 mM Tris pH7.5, 150 mM NaCl, 1 mM EDTA, 1% (v/v) IGEPAL CA-630 (Sigma), 10% (v/v) glycerol, supplemented with protease inhibitor cocktail (Sigma)) and transferred to 1.5 ml centrifugation tubes, spun down for 10 minutes at ∼20 000g at 4°C and the supernatants were transferred to new tubes. Total protein concentration was measured by BCA (Thermo) and 15 μg of protein were denatured by boiling the samples at 100 °C for 5 minutes in Laemmli buffer (50 mM Tris-HCl (pH 6.8), 2% SDS, 10% glycerol, 0.025% bromophenol blue, and 0.1 M β-mercaptoethanol). Samples were subjected to 12% SDS-PAGE followed by western blot onto PVDF membranes. Endogenous Gb1-4 subunits, HA-KCTD5 and b-tubulin were detected with anti-pan Gb (1/2000, #610288, BD Transduction laboratories), anti-HA (1/2000, #901501, BioLegend) and anti-b-tubulin (1/20000, #32-2600, Invitrogen) antibodies respectively, diluted in TBST (Tris-buffer saline with 0.1% Tween-20) containing 3% non-fat dry milk overnight at 4 °C. Membranes were washed thrice with TBST followed by one hour incubation with anti-mouse conjugated to HRP at RT followed by 3 TBST washes. Bands were revealed using ECL (ECL Select from GE). Intensity was quantified using ImageJ (https://imagej.nih.gov/ij/). Statistical analyses were performed using GraphPad Prism9.

### Ubiquitylation of Gbg *in cellulo*

Parental and KCTD5 KO cells were transfected with pcDNA3.1, TAP-HA-Gb1, HA-Gg2, and Flag-ubiquitin. Two days after transfection, cells were treated with 10 mM MG132 in DMSO for 8 h at 37 °C. Control cells were treated with DMSO. Treated cells were placed on ice, washed once with ice-cold PBS, and resuspended in 1 ml lysis buffer (50 mM Hepes-NaOH, pH 8.0, 150 mM NaCl, 0.3% Igepal CA-630, 2 mM EDTA, 2 mM DTT, 15 mM NEM and a protein inhibitor cocktail (Sigma)). Cells were lysed for 30 min with gentle rotation at 4 °C. Lysates were cleared by centrifugation at 21 300 × g for 10 min. Protein concentrations were measured using the Bradford protein assay (Bio-Rad, Mississauga, Ontario, Canada) using BSA as standard. For streptavidin purification, streptavidin magnetic sepharose beads (Cytiva, Uppsala, Sweden) were washed twice and incubated with 500 μg of total protein lysate for 3 h with gentle rotation at 4 °C. Beads were washed thrice with lysis buffer and proteins eluted with Laemmli sample buffer (62.5 mM Tris–HCl pH 6.8, 16.25% glycerol, 2% SDS, 5% β-mercaptoethanol, 0.025% bromophenol blue) and heated to 65 °C for 15 min followed by SDS-PAGE and western blot. Ubiquitinated proteins were detected using an anti-FLAG antibody (F7425, Sigma), HA-TAP-Gb1 and endogenous Gb1-4 were detected with anti-HA (16B112, BioLegend) and anti-Gb1-4 (sc-378, Santa Cruz) antibodies respectively.

### Detecting KCTD5/Gβ1 interactions

A *T*andem *A*ffinity *P*urification (TAP)-HA-Gβ1 vector was generated by PCR amplification of human Gβ1 from pcDNA3.1(+)-Flag-Gβ1 (cDNA Resource Center) using forward 5’- GTCCGAATTCATGAGTGAGCTTGACCAGTTAC –3’ and reverse 5’- GCTGGATCCTTAGTTCCAGATCTTGAG -3’ primers. Amplified HA-tagged Gβ1 cDNA was inserted at EcoRI and BamHI sites in pIRESpuro-GLUE-N1 (69). TAP-HA-Smad2 was similarly constructed, where HA-tagged human Smad2 was amplified using forward 5’- CCTAGGAATTCCAATCGTCCATCTTGCC -3’, and reverse 5’-GCAGTCTGCAGTTATGACATGCTTGAGC -3’ primers. The fragment was inserted at EcoRI and PstI restriction sites in pIRESpuro-GLUE-N1. KCTD5-Flag was subsequently generated, where KCTD5-Flag was amplified by PCR and ligated into a BamHI and EcoRI digested pcDNA3.1(+). TAP-HA-Gβ1 or TAP-HA-Smad2 stable lines were transfected with HA-KCTD5. Forty-eight hours after transfection, cells were lysed in TAP lysis buffer (10% glycerol, 50 mM HEPES NaOH pH 8.0, 2 mM EDTA, 0.3% Igepal CA-630, 2 mM DTT, 10 mM NaF, 0.25 mM sodium orthovanadate, 50 mM β-glycerolphosphate, protease inhibitor cocktail) at 4°C for 30 minutes with gentle agitation. Following a freeze-thaw cycle to increase protein recovery, samples were centrifuged for 20 minutes at 12 000g. Streptavidin purification was performed as described above. Samples were washed three times in the TAP-lysis buffer and following complete aspiration, beads were resuspended in 150 μl of elution buffer (TAP-lysis buffer containing 10 mM D-Biotin, pH 7.5-8.0), and left for 5 minutes on ice with mixing by flicking. The last step was repeated once, and eluates combined followed by SDS-PAGE and western blot. Co-immunoprecipitation was performed similarly. HEK 293 cells stably expressing TAP-HA-Gβ1 were transfected with KCTD5-FLAG. Western blot analysis of the samples were done using anti-HA (1:2 000; BioLegend) and anti-Flag M2 (1:5 000; Sigma) as primary antibodies.

## Data Availability

The cryo-EM maps and associated atomic models of top and bottom complex and conformational states A, B, C and D have been deposited in the Electron Microscopy Data Bank with accession codes EMD-41994, EMD-41995, EMD-41996, EMD-42000, EMD-42004, and EMD-42008 and the Protein Data Bank with accession codes 8U7Z, 8U80, 8U81, 8U82, 8U83 and 8U84, respectively. Maps and models from the multi-body analysis are available from Zenodo at https://doi.org/10.5281/zenodo.8341597.

## Acknowledgments

This research was funded by a Canadian Institutes of Health Research grant PJT162186 to GGP. DMN was supported by Canadian Institutes of Health Research CGS-M. We thank Frank Sicheri, Samuel Lunenfeld Institute, for the expression vector for ARIH1 and the SGC for expression vectors for UBE1 and UBE2L3. TEH is supported by funding from the Heart and Stroke Society of Canada and PJT159687 from CIHR. TEH holds the Canadian Pacific Chair in Biotechnology. We thank Tiffany Villleneuve and Fenneke KleinJan for instrument support, Dr. Richard Wargachuk for construction of KCTD5-FLAG and Dr. Tamara Ouspenskaia for construction of TAP-Smad2. Titan Krios cryo-EM data were collected at the Toronto High-Resolution High-Throughput Cryo-EM facility supported by the Canada Foundation for Innovation and Ontario Research Fund.

## Notes

### Competing Interest Statement

The authors have declared no competing interest.

## References

1. D.-V. Rusnac, N. Zheng, Structural Biology of CRL Ubiquitin Ligases. Adv. Exp. Med. Biol. 1217, 9–31 (2020).

2. J. W. Harper, B. A. Schulman, Cullin-RING Ubiquitin Ligase Regulatory Circuits: A Quarter Century Beyond the F-Box Hypothesis. Annu. Rev. Biochem. 90, 403–429 (2021).

3. P. Wang, J. Song, D. Ye, CRL3s: The BTB-CUL3-RING E3 Ubiquitin Ligases. Adv. Exp. Med. Biol. 1217, 211–223 (2020).

4. W. J. Errington, M. Q. Khan, S. A. Bueler, J. L. Rubinstein, A. Chakrabartty, G. G. Privé, Adaptor protein self-assembly drives the control of a cullin-RING ubiquitin ligase. Structure 20, 1141–1153 (2012).

5. P. Canning, C. D. O. Cooper, T. Krojer, J. W. Murray, A. C. W. Pike, A. Chaikuad, T. Keates, C. Thangaratnarajah, V. Hojzan, B. D. Marsden, O. Gileadi, S. Knapp, F. von Delft, A. N. Bullock, Structural basis for Cul3 protein assembly with the BTB-Kelch family of E3 ubiquitin ligases. J. Biol. Chem. 288, 7803–7814 (2013).

6. A. X. Ji, A. Chu, T. K. Nielsen, S. Benlekbir, J. L. Rubinstein, G. G. Privé, Structural Insights into KCTD Protein Assembly and Cullin3 Recognition. J. Mol. Biol. 428, 92–107 (2016).

7. G. Smaldone, L. Pirone, E. Pedone, T. Marlovits, L. Vitagliano, L. Ciccarelli, The BTB domains of the potassium channel tetramerization domain proteins prevalently assume pentameric states. FEBS Lett. 590, 1663–1671 (2016).

8. D. M. Pinkas, C. E. Sanvitale, J. C. Bufton, F. J. Sorrell, N. Solcan, R. Chalk, J. Doutch, A. N. Bullock, Structural complexity in the KCTD family of Cullin3-dependent E3 ubiquitin ligases. Biochem. J. 474, 3747–3761 (2017).

9. M. J. Cuneo, B. G. O’Flynn, Y.-H. Lo, N. Sabri, T. Mittag, Higher-order SPOP assembly reveals a basis for cancer mutant dysregulation. Mol. Cell 83, 731–745.e4 (2023).

10. S. Toma-Fukai, T. Shimizu, Structural Diversity of Ubiquitin E3 Ligase. Molecules 26, 6682 (2021).

11. X. Tang, S. Orlicky, Z. Lin, A. Willems, D. Neculai, D. Ceccarelli, F. Mercurio, B. H. Shilton, F. Sicheri, M. Tyers, Suprafacial orientation of the SCFCdc4 dimer accommodates multiple geometries for substrate ubiquitination. Cell 129, 1165–1176 (2007).

12. B. Hao, S. Oehlmann, M. E. Sowa, J. W. Harper, N. P. Pavletich, Structure of a Fbw7- Skp1-cyclin E complex: multisite-phosphorylated substrate recognition by SCF ubiquitin ligases. Mol. Cell 26, 131–143 (2007).

13. S. M. Ahmed, A. M. Daulat, A. Meunier, S. Angers, G protein betagamma subunits regulate cell adhesion through Rap1a and its effector Radil. J. Biol. Chem. 285, 6538–6551 (2010).

14. R. Campden, D. Pétrin, M. Robitaille, N. Audet, S. Gora, S. Angers, T. E. Hébert, Tandem affinity purification to identify cytosolic and nuclear gβγ-interacting proteins. Methods Mol. Biol. Clifton NJ 1234, 161–184 (2015).

15. Z. Zha, X. Han, M. D. Smith, Y. Liu, P. M. Giguère, D. Kopanja, P. Raychaudhuri, D. P. Siderovski, K.-L. Guan, Q.-Y. Lei, Y. Xiong, A Non-Canonical Function of Gβ as a Subunit of E3 Ligase in Targeting GRK2 Ubiquitylation. Mol. Cell 58, 794–803 (2015).

16. M. Brockmann, V. A. Blomen, J. Nieuwenhuis, E. Stickel, M. Raaben, O. B. Bleijerveld, A. F. M. Altelaar, L. T. Jae, T. R. Brummelkamp, Genetic wiring maps of single-cell protein states reveal an off-switch for GPCR signalling. Nature 546, 307–311 (2017).

17. B. D. Young, J. Sha, A. A. Vashisht, J. A. Wohlschlegel, Human Multisubunit E3 Ubiquitin Ligase Required for Heterotrimeric G-Protein β-Subunit Ubiquitination and Downstream Signaling. J. Proteome Res. 20, 4318–4330 (2021).

18. B. S. Muntean, S. Marwari, X. Li, D. C. Sloan, B. D. Young, J. A. Wohlschlegel, K. A. Martemyanov, Members of the KCTD family are major regulators of cAMP signaling. Proc. Natl. Acad. Sci. U. S. A. 119, e2119237119 (2022).

19. D. C. Sloan, C. E. Cryan, B. S. Muntean, Multiple potassium channel tetramerization domain (KCTD) family members interact with Gβγ, with effects on cAMP signaling. J. Biol. Chem. 299, 102924 (2023).

20. W. Jiang, W. Wang, Y. Kong, S. Zheng, Structural basis for the ubiquitination of G protein βγ subunits by KCTD5/Cullin3 E3 ligase. Sci. Adv. 9, eadg8369 (2023).

21. Q. Li, D. A. Kellner, H. A. M. Hatch, T. Yumita, S. Sanchez, R. P. Machold, C. A. Frank, N. Stavropoulos, Conserved properties of Drosophila Insomniac link sleep regulation and synaptic function. PLoS Genet. 13, e1006815 (2017).

22. R. Barfield, et al., Epigenome-wide association analysis of daytime sleepiness in the Multi-Ethnic Study of Atherosclerosis reveals African-American-specific associations. Sleep 42, zsz101 (2019).

23. Z. Liu, Y. Xiang, G. Sun, The KCTD family of proteins: structure, function, disease relevance. Cell Biosci. 3, 45 (2013).

24. M. Skoblov, A. Marakhonov, E. Marakasova, A. Guskova, V. Chandhoke, A. Birerdinc, A. Baranova, Protein partners of KCTD proteins provide insights about their functional roles in cell differentiation and vertebrate development. BioEssays News Rev. Mol. Cell. Dev. Biol. 35, 586–596 (2013).

25. X. Teng, A. Aouacheria, L. Lionnard, K. A. Metz, L. Soane, A. Kamiya, J. M. Hardwick, KCTD: A new gene family involved in neurodevelopmental and neuropsychiatric disorders. CNS Neurosci. Ther. 25, 887–902 (2019).

26. A. Angrisani, A. Di Fiore, E. De Smaele, M. Moretti, The emerging role of the KCTD proteins in cancer. Cell Commun. Signal. CCS 19, 56 (2021).

27. M. Tennakoon, K. Senarath, D. Kankanamge, K. Ratnayake, D. Wijayaratna, K. Olupothage, S. Ubeysinghe, K. Martins-Cannavino, T. E. Hébert, A. Karunarathne, Subtype-dependent regulation of Gβγ signalling. Cell. Signal. 82, 109947 (2021).

28. S. M. Khan, R. Sleno, S. Gora, P. Zylbergold, J.-P. Laverdure, J.-C. Labbé, G. J. Miller, T. E. Hébert, The expanding roles of Gβγ subunits in G protein-coupled receptor signaling and drug action. Pharmacol. Rev. 65, 545–577 (2013).

29. S. M. Khan, J. Y. Sung, T. E. Hébert, Gβγ subunits-Different spaces, different faces. Pharmacol. Res. 111, 434–441 (2016).

30. K. Baek, D. T. Krist, J. R. Prabu, S. Hill, M. Klügel, L.-M. Neumaier, S. von Gronau, G. Kleiger, B. A. Schulman, NEDD8 nucleates a multivalent cullin-RING-UBE2D ubiquitin ligation assembly. Nature 578, 461–466 (2020).

31. D. Horn-Ghetko, D. T. Krist, J. R. Prabu, K. Baek, M. P. C. Mulder, M. Klügel, D. C. Scott, H. Ovaa, G. Kleiger, B. A. Schulman, Ubiquitin ligation to F-box protein targets by SCF-RBR E3-E3 super-assembly. Nature 590, 671–676 (2021).

32. E. L. Mena, P. Jevtić, B. J. Greber, C. L. Gee, B. G. Lew, D. Akopian, E. Nogales, J. Kuriyan, M. Rape, Structural basis for dimerization quality control. Nature 586, 452–456 (2020).

33. M. Jenkyn-Bedford, M. L. Jones, Y. Baris, K. P. M. Labib, G. Cannone, J. T. P. Yeeles, T. D. Deegan, A conserved mechanism for regulating replisome disassembly in eukaryotes. Nature 600, 743–747 (2021).

34. D. C. Scott, D. Y. Rhee, D. M. Duda, I. R. Kelsall, J. L. Olszewski, J. A. Paulo, A. de Jong, H. Ovaa, A. F. Alpi, J. W. Harper, B. A. Schulman, Two Distinct Types of E3 Ligases Work in Unison to Regulate Substrate Ubiquitylation. Cell 166, 1198–1214.e24 (2016).

35. N. Balasco, L. Pirone, G. Smaldone, S. Di Gaetano, L. Esposito, E. M. Pedone, L. Vitagliano, Molecular recognition of Cullin3 by KCTDs: insights from experimental and computational investigations. Biochim. Biophys. Acta 1844, 1289–1298 (2014).

36. I. S. Dementieva, V. Tereshko, Z. A. McCrossan, E. Solomaha, D. Araki, C. Xu, N. Grigorieff, S. A. N. Goldstein, Pentameric assembly of potassium channel tetramerization domain-containing protein 5. J. Mol. Biol. 387, 175–191 (2009).

37. K. Wu, J. Kovacev, Z.-Q. Pan, Priming and extending: a UbcH5/Cdc34 E2 handoff mechanism for polyubiquitination on a SCF substrate. Mol. Cell 37, 784–796 (2010).

38. S. Hill, K. Reichermeier, D. C. Scott, L. Samentar, J. Coulombe-Huntington, L. Izzi, X. Tang, R. Ibarra, T. Bertomeu, A. Moradian, M. J. Sweredoski, N. Caberoy, B. A. Schulman, F. Sicheri, M. Tyers, G. Kleiger, Robust cullin-RING ligase function is established by a multiplicity of poly-ubiquitylation pathways. eLife 8, e51163 (2019).

39. D. Barone, N. Balasco, L. Vitagliano, KCTD5 is endowed with large, functionally relevant, interdomain motions. J. Biomol. Struct. Dyn. 34, 1725–1735 (2016).

40. M. A. Wall, D. E. Coleman, E. Lee, J. A. Iñiguez-Lluhi, B. A. Posner, A. G. Gilman, S. R. Sprang, The structure of the G protein heterotrimer Gi alpha 1 beta 1 gamma 2. Cell 83, 1047–1058 (1995).

41. D. T. Lodowski, J. A. Pitcher, W. D. Capel, R. J. Lefkowitz, J. J. G. Tesmer, Keeping G proteins at bay: a complex between G protein-coupled receptor kinase 2 and Gbetagamma. Science 300, 1256–1262 (2003).

42. S. Zheng, N. Abreu, J. Levitz, A. C. Kruse, Structural basis for KCTD-mediated rapid desensitization of GABAB signalling. Nature 567, 127–131 (2019).

43. A. V. Smrcka, I. Fisher, G-protein βγ subunits as multi-functional scaffolds and transducers in G-protein-coupled receptor signaling. Cell. Mol. Life Sci. CMLS 76, 4447–4459 (2019).

44. A. Punjani, D. J. Fleet, 3DFlex: determining structure and motion of flexible proteins from cryo-EM. Nat. Methods 20, 860–870 (2023).

45. P. J. Stogios, G. S. Downs, J. J. S. Jauhal, S. K. Nandra, G. G. Privé, Sequence and structural analysis of BTB domain proteins. Genome Biol. 6, R82 (2005).

46. K. Baek, D. C. Scott, L. T. Henneberg, M. T. King, M. Mann, B. A. Schulman, Systemwide disassembly and assembly of SCF ubiquitin ligase complexes. Cell 186, 2492 (2023).

47. M. Shaaban, J. A. Clapperton, S. Ding, S. Kunzelmann, M.-E. Mäeots, S. L. Maslen, J. M. Skehel, R. I. Enchev, Structural and mechanistic insights into the CAND1-mediated SCF substrate receptor exchange. Mol. Cell 83, 2332–2346.e8 (2023).

48. A. X. Ji, G. G. Privé, Crystal structure of KLHL3 in complex with Cullin3. PloS One 8, e60445 (2013).

49. T. Nakane, D. Kimanius, E. Lindahl, S. H. Scheres, Characterisation of molecular motions in cryo-EM single-particle data by multi-body refinement in RELION. eLife 7 (2018).

50. E. D. Zhong, T. Bepler, B. Berger, J. H. Davis, CryoDRGN: reconstruction of heterogeneous cryo-EM structures using neural networks. Nat. Methods 18, 176–185 (2021).

51. T. Mittag, S. Orlicky, W.-Y. Choy, X. Tang, H. Lin, F. Sicheri, L. E. Kay, M. Tyers, J. D. Forman-Kay, Dynamic equilibrium engagement of a polyvalent ligand with a single-site receptor. Proc. Natl. Acad. Sci. U. S. A. 105, 17772–17777 (2008).

52. K. Baek, D. C. Scott, B. A. Schulman, NEDD8 and ubiquitin ligation by cullin-RING E3 ligases. Curr. Opin. Struct. Biol. 67, 101–109 (2021).

53. N. W. Pierce, J. E. Lee, X. Liu, M. J. Sweredoski, R. L. J. Graham, E. A. Larimore, M. Rome, N. Zheng, B. E. Clurman, S. Hess, S. Shan, R. J. Deshaies, Cand1 promotes assembly of new SCF complexes through dynamic exchange of F box proteins. Cell 153, 206–215 (2013).

54. K. Martins-Cannavino, T. E. Hébert, Gβγ signaling from an eponymous past to a specific future. Cell Syst. 12, 289–290 (2021).

55. Z. Li, I. P. Michael, D. Zhou, A. Nagy, J. M. Rini, Simple piggyBac transposon-based mammalian cell expression system for inducible protein production. Proc. Natl. Acad. Sci. U. S. A. 110, 5004–5009 (2013).

56. C. R. Marr, S. Benlekbir, J. L. Rubinstein, Fabrication of carbon films with ∼ 500nm holes for cryo-EM with a direct detector device. J. Struct. Biol. 185, 42–47 (2014).

57. A. Punjani, J. L. Rubinstein, D. J. Fleet, M. A. Brubaker, cryoSPARC: algorithms for rapid unsupervised cryo-EM structure determination. Nat. Methods 14, 290–296 (2017).

58. A. Punjani, H. Zhang, D. J. Fleet, Non-uniform refinement: adaptive regularization improves single-particle cryo-EM reconstruction. Nat. Methods 17, 1214–1221 (2020).

59. R. Sanchez-Garcia, J. Gomez-Blanco, A. Cuervo, J. M. Carazo, C. O. S. Sorzano, J. Vargas, DeepEMhancer: a deep learning solution for cryo-EM volume post-processing. *Commun*. Biol. 4, 874 (2021).

60. P. Emsley, B. Lohkamp, W. G. Scott, K. Cowtan, Features and development of Coot. Acta Crystallogr. D Biol. Crystallogr. 66, 486–501 (2010).

61. P. D. Adams, P. V. Afonine, G. Bunkóczi, V. B. Chen, I. W. Davis, N. Echols, J. J. Headd, L.-W. Hung, G. J. Kapral, R. W. Grosse-Kunstleve, A. J. McCoy, N. W. Moriarty, R. Oeffner, R. J. Read, D. C. Richardson, J. S. Richardson, T. C. Terwilliger, P. H. Zwart, PHENIX: a comprehensive Python-based system for macromolecular structure solution. Acta Crystallogr. D Biol. Crystallogr. 66, 213–221 (2010).

62. D. Kimanius, L. Dong, G. Sharov, T. Nakane, S. H. W. Scheres, New tools for automated cryo-EM single-particle analysis in RELION-4.0. Biochem. J. 478, 4169–4185 (2021).

63. T. Nakane, S. H. W. Scheres, Multi-body Refinement of Cryo-EM Images in RELION. Methods Mol. Biol. Clifton NJ 2215, 145–160 (2021).

64. R. T. Kidmose, J. Juhl, P. Nissen, T. Boesen, J. L. Karlsen, B. P. Pedersen, Namdinator - automatic molecular dynamics flexible fitting of structural models into cryo-EM and crystallography experimental maps. IUCrJ 6, 526–531 (2019).

65. E. F. Pettersen, T. D. Goddard, C. C. Huang, G. S. Couch, D. M. Greenblatt, E. C. Meng, T. E. Ferrin, UCSF Chimera--a visualization system for exploratory research and analysis. J. Comput. Chem. 25, 1605–1612 (2004).

66. *The PyMOL Molecular Graphics System* (Schrödinger, LLC).

67. A. Morin, B. Eisenbraun, J. Key, P. C. Sanschagrin, M. A. Timony, M. Ottaviano, P. Sliz, Collaboration gets the most out of software. eLife 2, e01456 (2013).

68. R. V. Rebois, M. Robitaille, C. Galés, D. J. Dupré, A. Baragli, P. Trieu, N. Ethier, M. Bouvier, T. E. Hébert, Heterotrimeric G proteins form stable complexes with adenylyl cyclase and Kir3.1 channels in living cells. J. Cell Sci. 119, 2807–2818 (2006).

69. S. Angers, C. J. Thorpe, T. L. Biechele, S. J. Goldenberg, N. Zheng, M. J. MacCoss, R. T. Moon, The KLHL12-Cullin-3 ubiquitin ligase negatively regulates the Wnt-beta-catenin pathway by targeting Dishevelled for degradation. Nat. Cell Biol. 8, 348–357 (2006).

