## Supplemental Figures and Tables for "Structure and dynamics of a pentameric KCTD5/Cullin3/Gβγ E3 ubiquitin ligase complex"

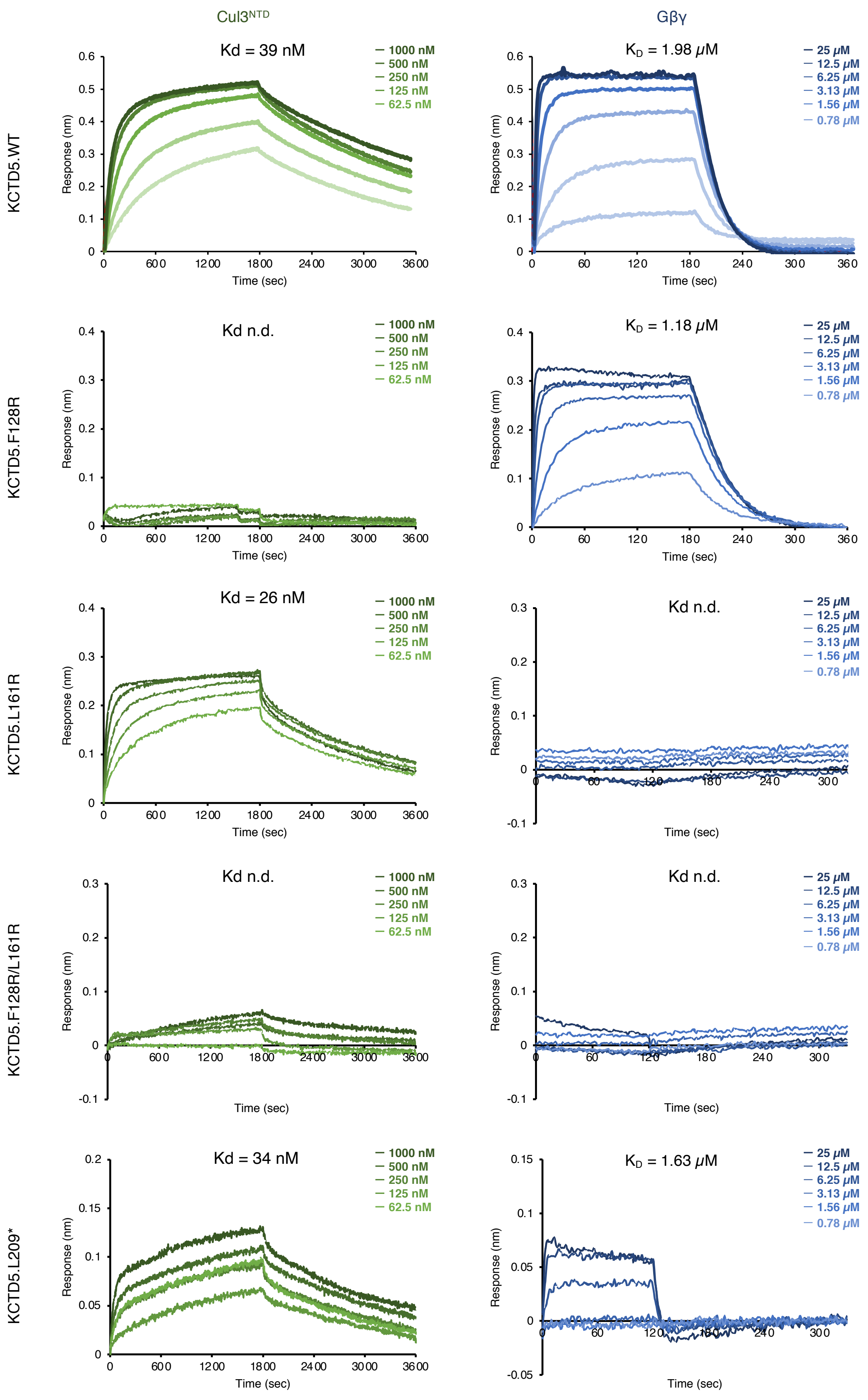


**Fig. S1.** BLI data of anchored KCTD5 tested with Cul3$^{NTD}$ (left) and Gβγ (right). n.d.: not determined.

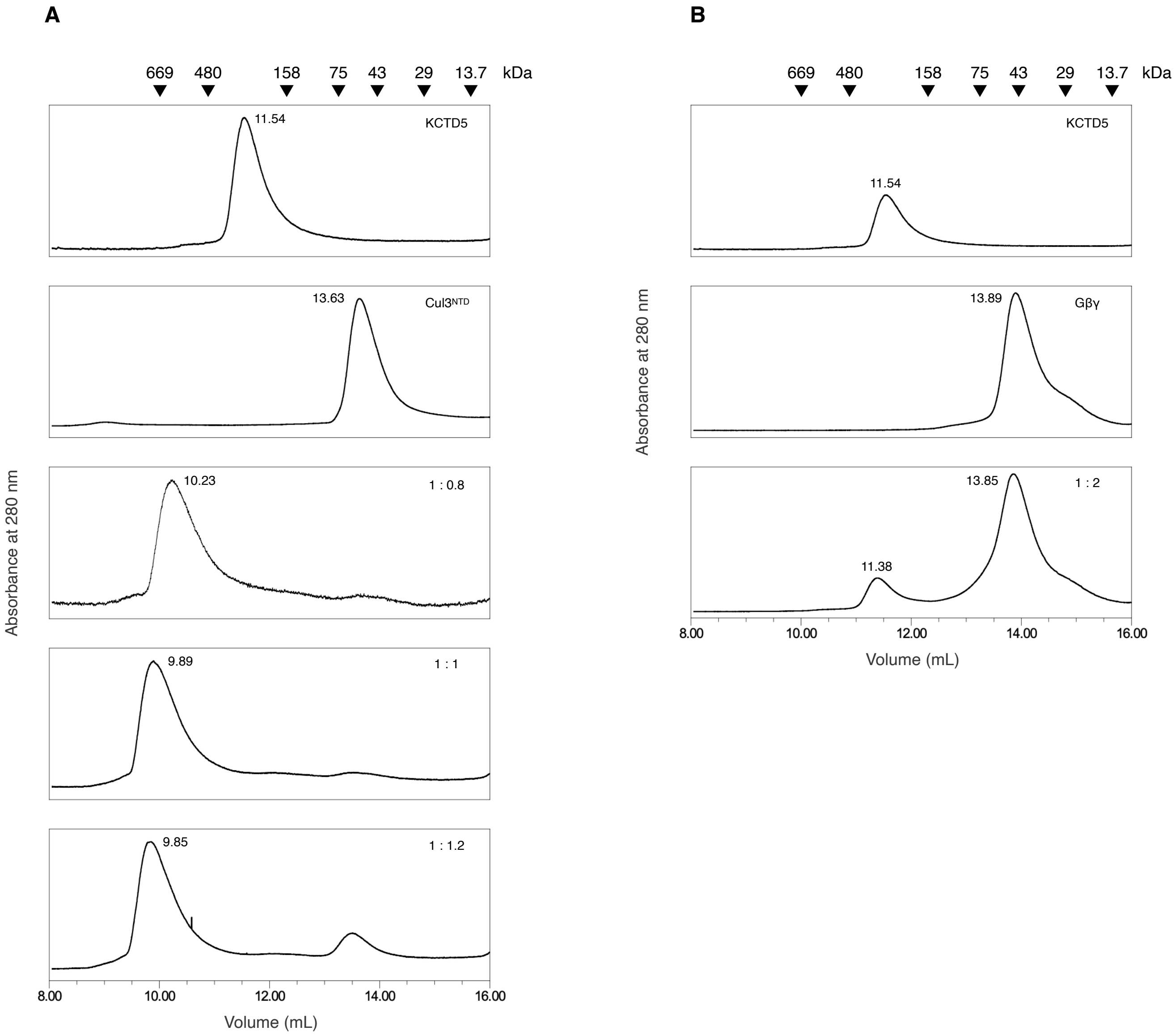


**Fig. S2.** SEC elution profiles. (A) KCTD5 and Cul3$^{NTD}$. The lower three panels are mixtures of the two proteins at increasing KCTD5:Cul3$^{NTD}$ molar ratios. (B) KCTD5 and Gβɣ. The third panel is a mixture with a 2-fold excess of Gβɣ. Molecular weight markers are shown above the profiles, and elution volumes are indicated near the peaks.

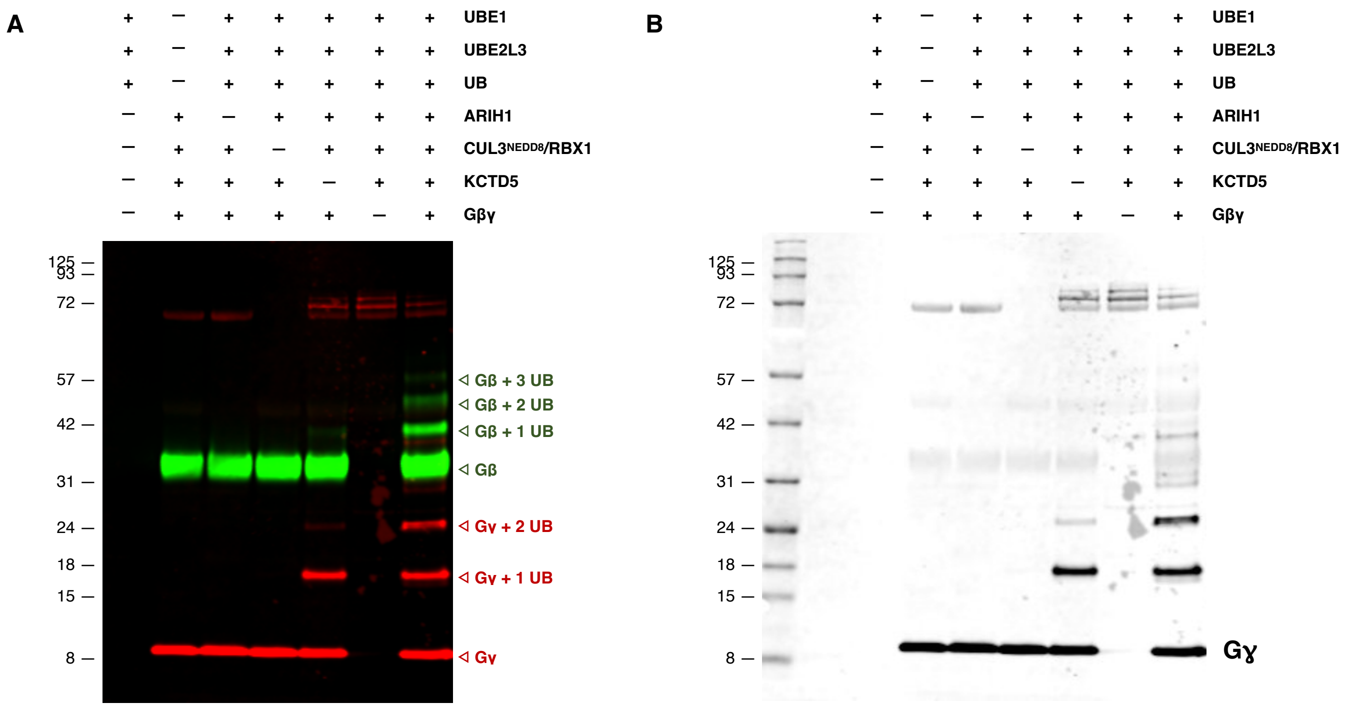


**Fig. S3.** Ubiquitylation of Gβɣ by CRL3$^{KCTD5}$. (A) Western blot image (anti-Gβ green, anti Gγ red). (B) Greyscale images of the Gγ channel. The greyscale image for the Gβ channel is shown in the main text Fig. 1.

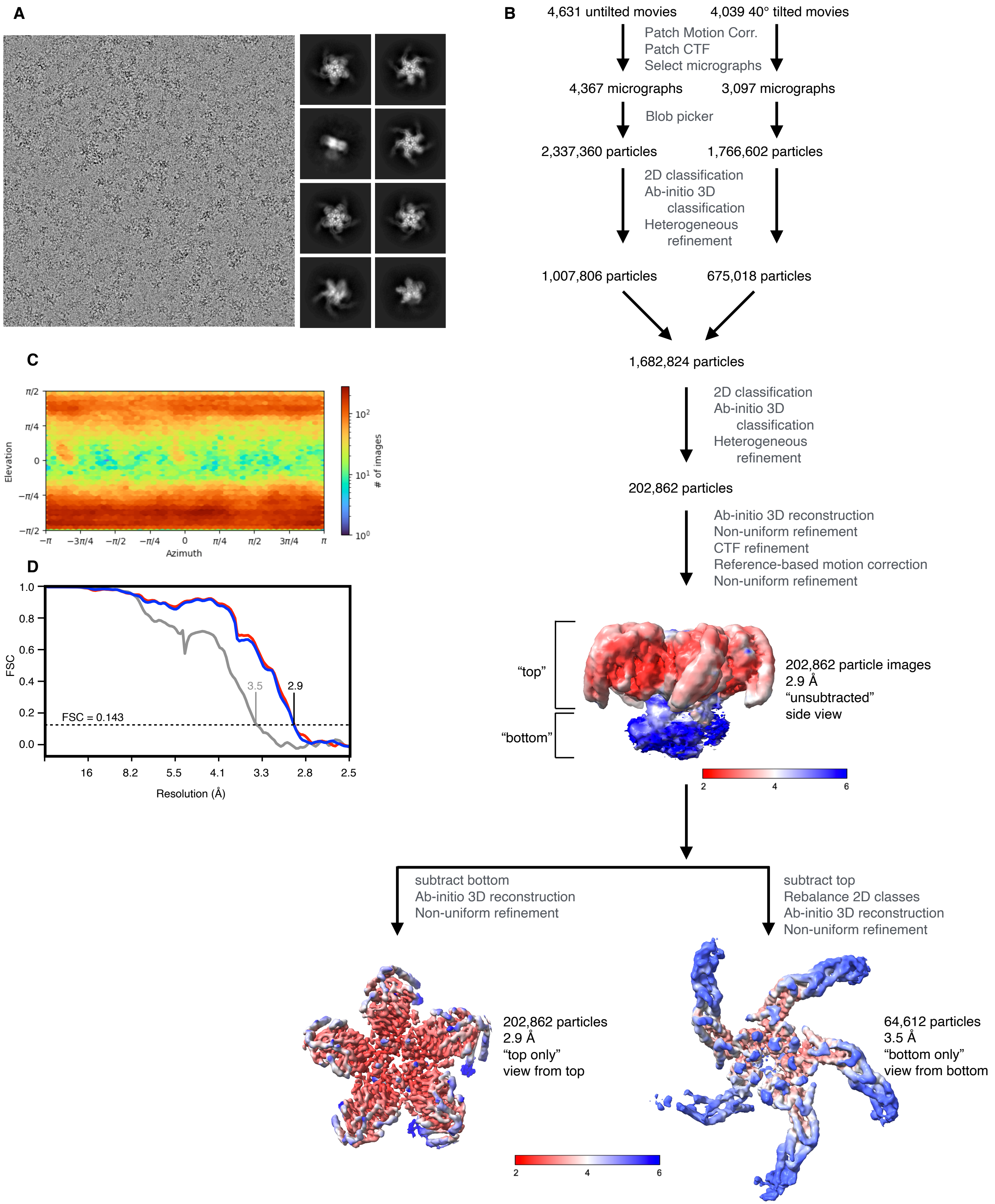


**Fig. S4.** Cryo-em of the KCTD5/Cul3$^{NTD}$/Gβγ complex (A) Representative micrograph (left) and 2D classes (right). (B) Cryo-em workflow leading to the unsubtracted, top only and bottom only maps. Maps are colored by local resolution. (C) Particle viewing direction distribution plot. (D) Fourier Shell Correlation (FSC) curves (corrected for the effects of masking) for the unsubtracted (blue), top only (red) and bottom only (grey) maps, respectively. Image analysis for the data shown in this figure was conducted with cryoSPARC.

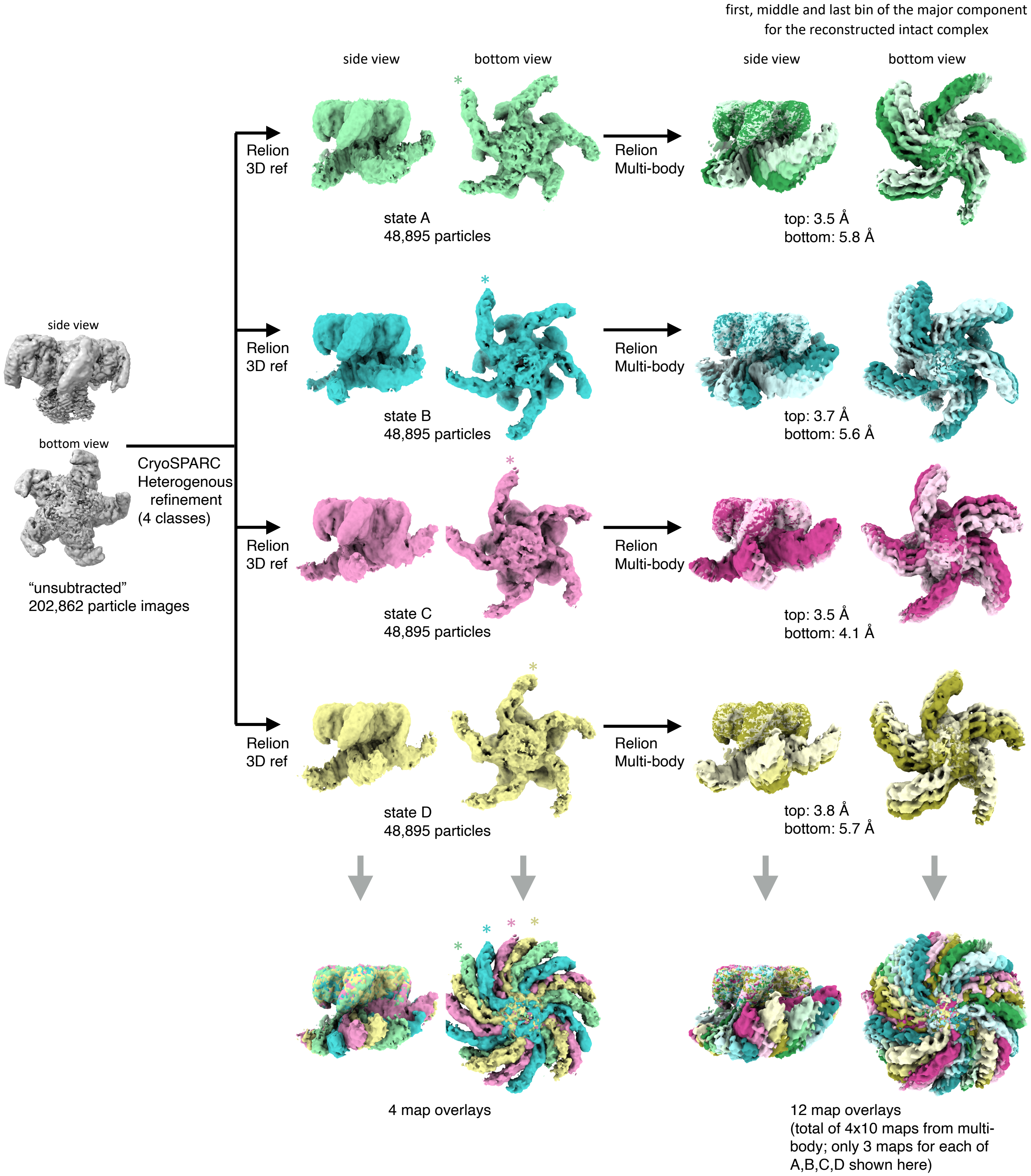


**Fig. S5.** Cryo-em maps of the intact complex. The particle image set was divided into subsets A, B, C and D via CryoSPARC heterogenous refinement, followed by 3D refinement in Relion. An asterisk indicates one Cul3$^{NTD}$ arm from the four Relion 3D refinements. The four states were then analyzed with Relion Multi-body refinement, each with two bodies: KCTD$^{CTD}$/Gβγ (top) and KCTD$^{BTB}$/Cul3$^{NTD}$ (bottom). Ten composite maps for one of the major component were calculated, and the 1st, 5th and 10th maps (bins) are shown at the right. All maps in this figure are are aligned to a common setting of the "top" KCTD5$^{CTD}$/Gbg moiety.

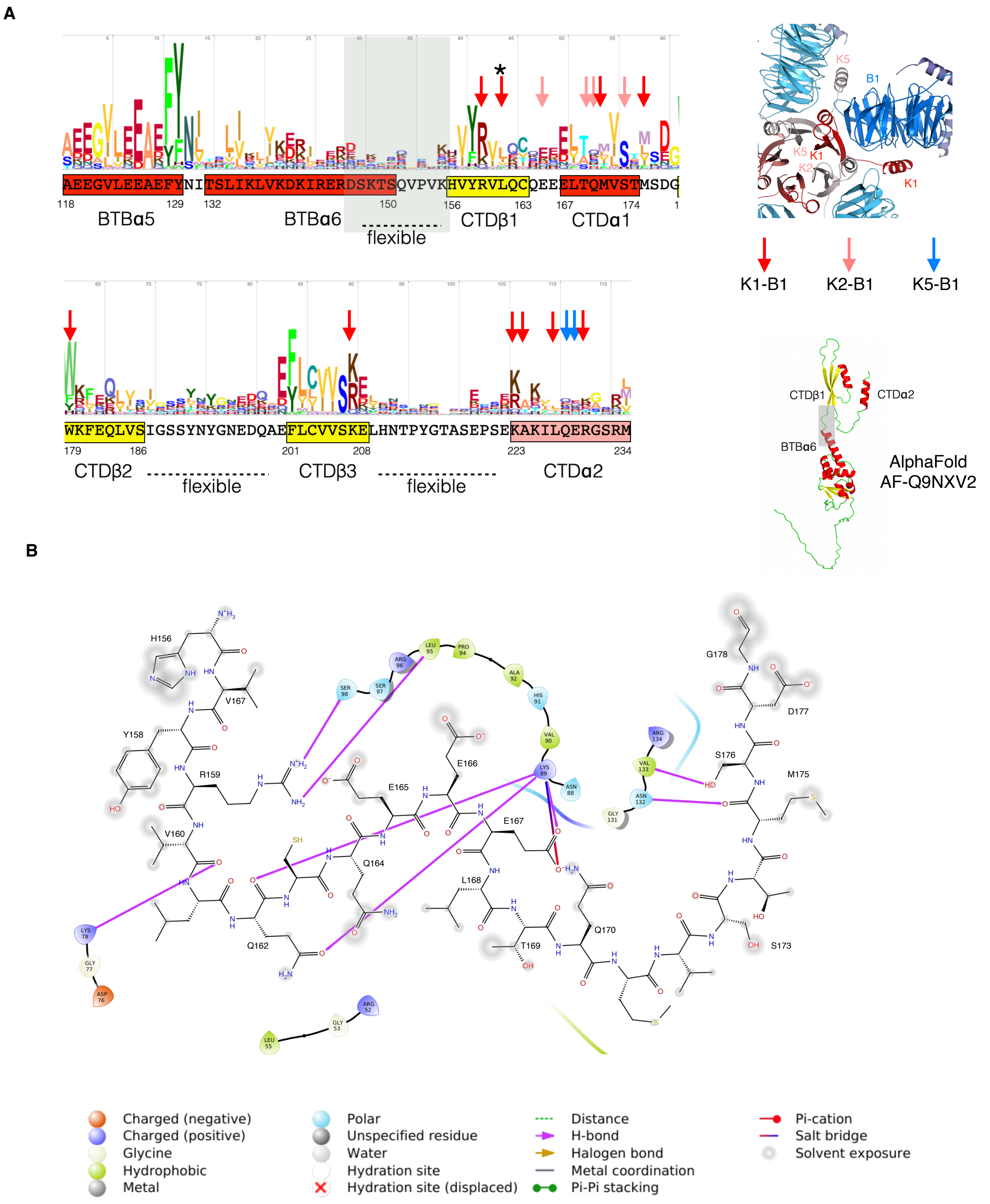


**Fig. S6** (A). Details of the KCTD5CTD/Gβγ interface. (A) Gβ contacts are shown along with sequence variability of KCTD5 orthologs. Arrows indicate major contacts according to they key on the right (K1, K2, K5 are KCTD5CTD chains and B1 is a Gβ chain). The asterisk indicates residue L161, mutated to L161R. The AlphaFold prediction for KCTD5 is shown on the right. (B) 2D representation of the contacts between KCTD5 chain K1 (sticks) and Gβ (spheres).

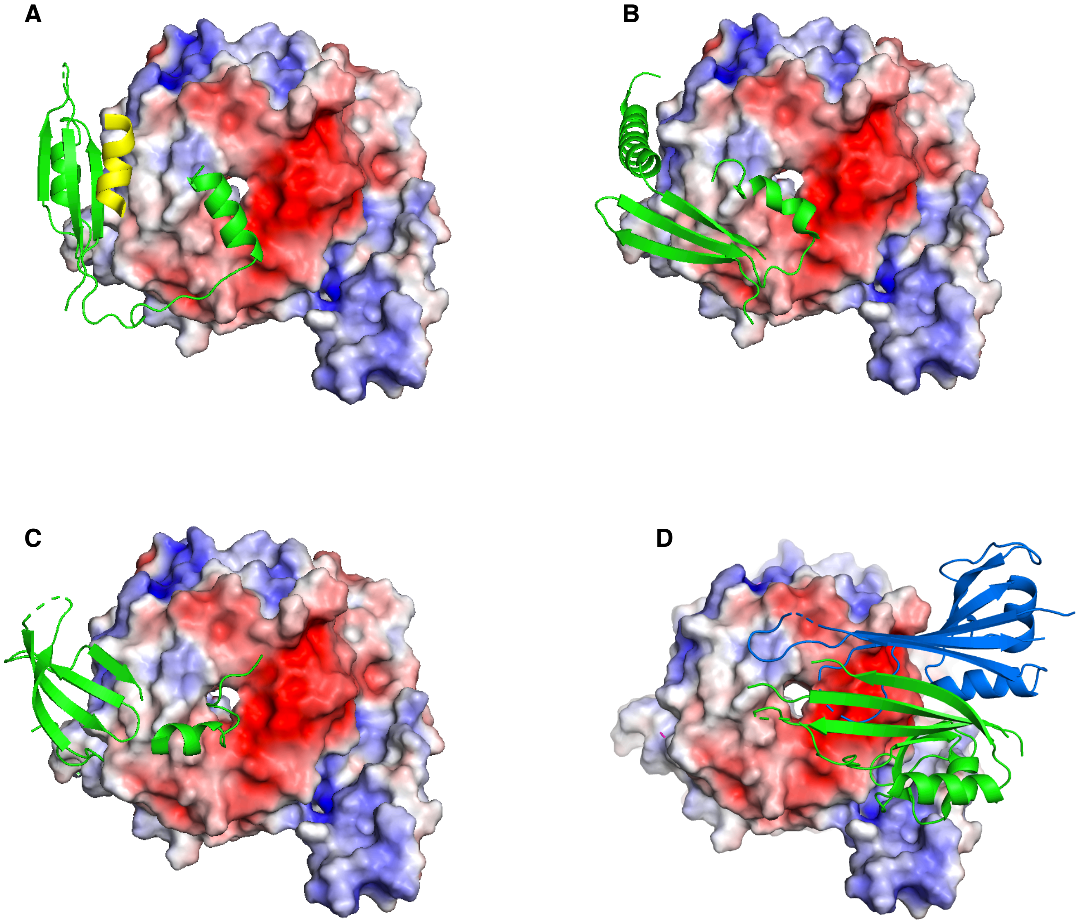


**Fig S7.** The electrostatic surface of Gβγ is shown in complex with (A) the primary (green) and secondary (yellow) subunits of KCTD5$^{CTD}$, (B) Gα (1GP2), (C) GRK2 (1OMW), and two adjacent KCTD12 H1 domains (green and blue) (6M8S).

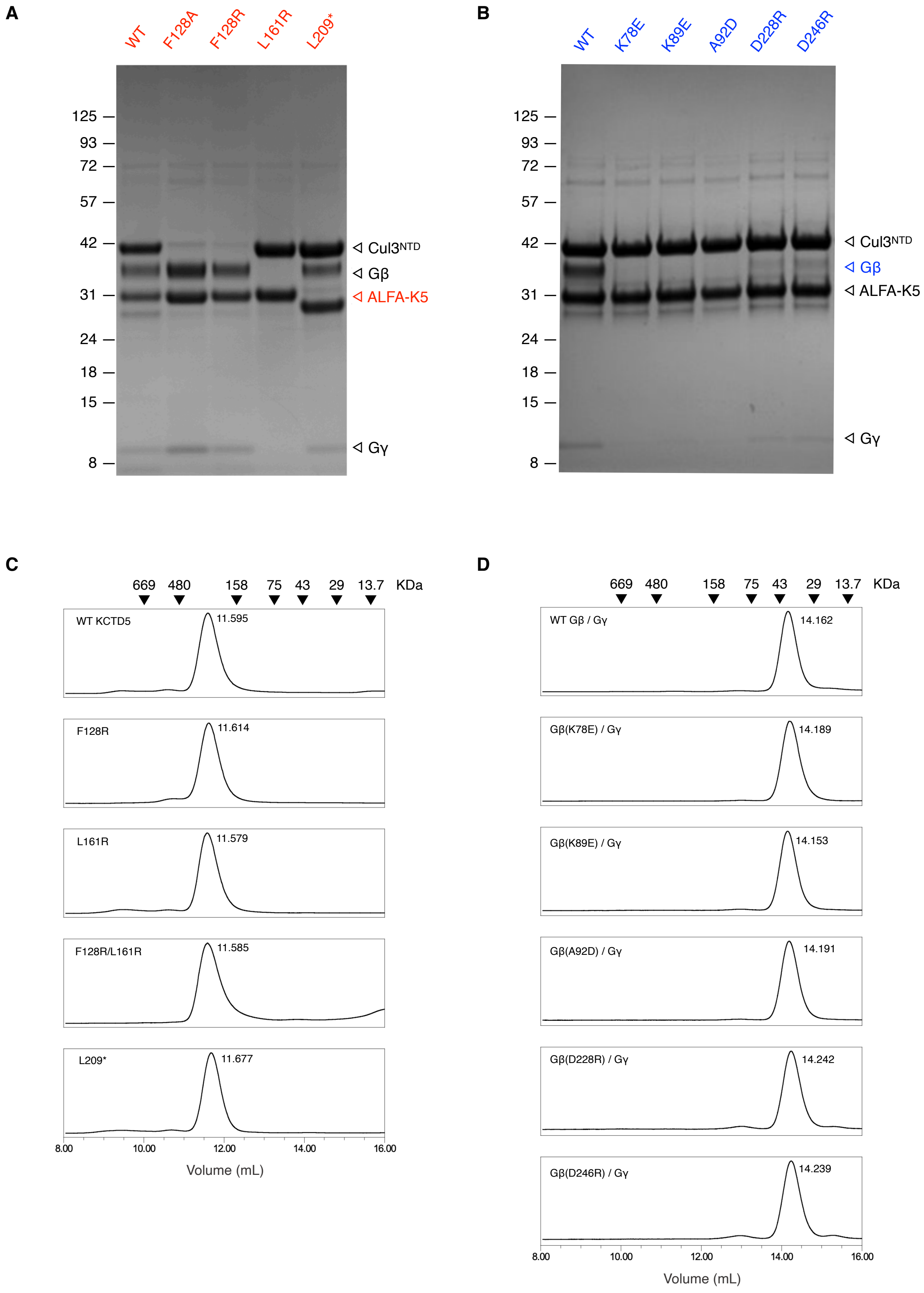


**Fig. S8.** Characterization of KCTD5 and Gβγ mutants. (A, B) ALFA-tagged KCTD5 pull down assay with $Cul3^{NTD}$ and Gβγ (Coomassie stained) with KCTD5 mutants and Gβ mutants. (C, D) SEC elution profiles for purified ALFA-KCTD5 and Gβγ. Molecular weight markers are shown above the SEC profiles, and elution volumes are indicated near the peaks.

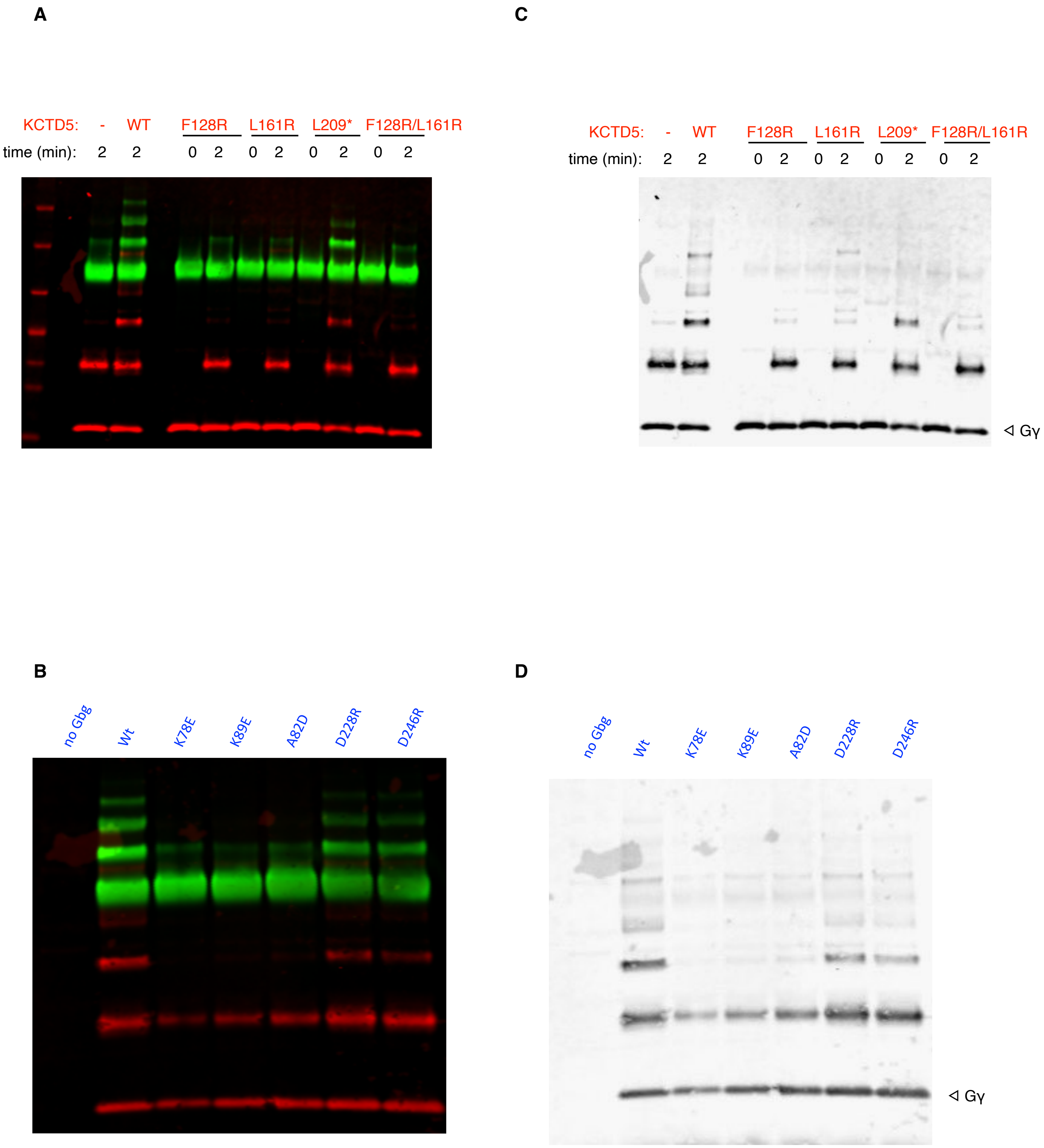


**Fig. S9.** Ubiquitylation activity of KCTD5 and Gβ mutants. (A, B) Western blot images (anti-Gβ green, anti Gγ red). (C,D) Greyscale images of the Gγ channel. The greyscale images for the Gβ channel is shown in the main text Fig. 2.

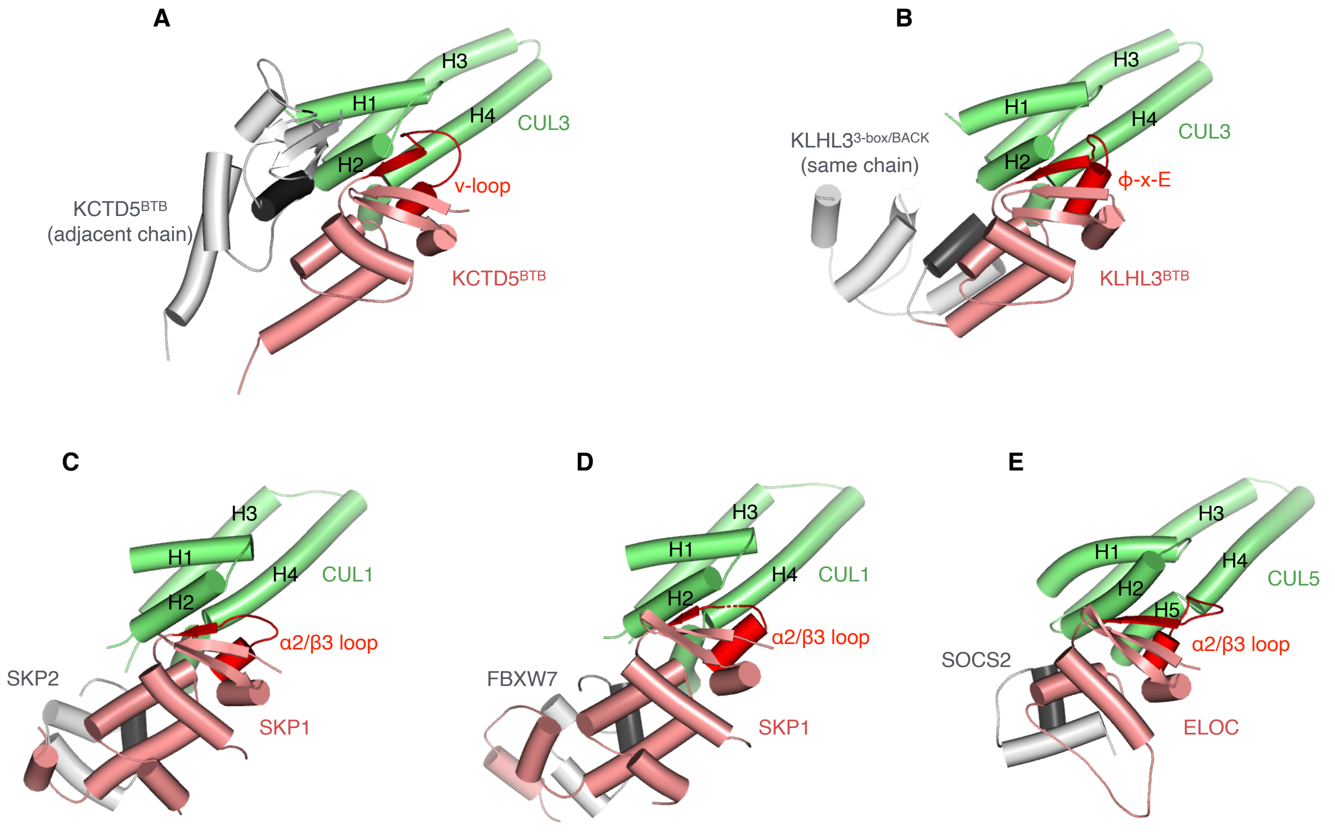


**Fig. S10.** Binding interactions in the cullin CR1 domain. In all panels, the cullin is green, BTB/SKP1/ELOC is pink with the α2/β3 region of the primary interface in red, and the distal domain is light grey with the contact helix in dark gray. (*A*) the KCTD5$^{BTB}$/CUL3 (this work), (*B*) KLHL3$^{BTB}$/CUL3 (4HXI), (*C*) SKP1/CUL1/SKP2 (1LDK), (*D*) SKP1/CUL1/FBXW7 (7B5M), (*E*) ELOC/CUL5/SOCS2 (4JGH).

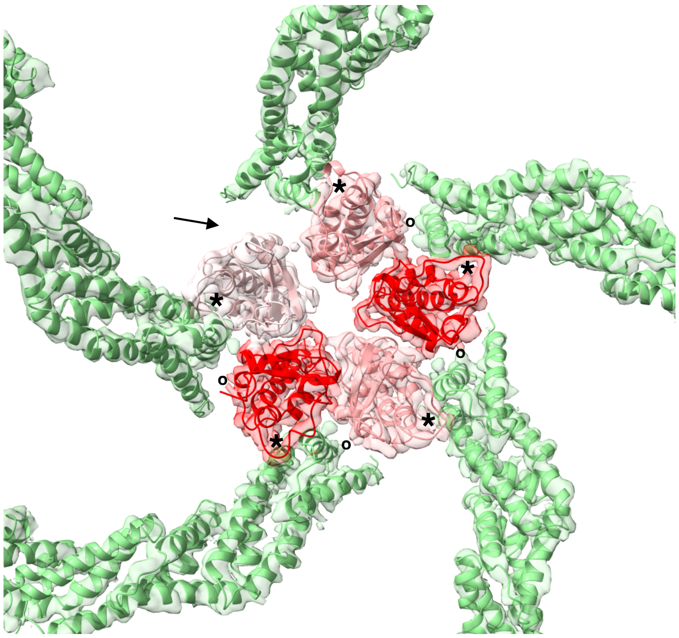

**Fig. S11.** Fit of the model into the bottom cryo-em map. Primary interfaces are indicated by "*" centered in the BTB v-loops, and distal interfaces are indicated by "o". The arrow indicates a position that lacks the distal interface due to BTB ring opening.

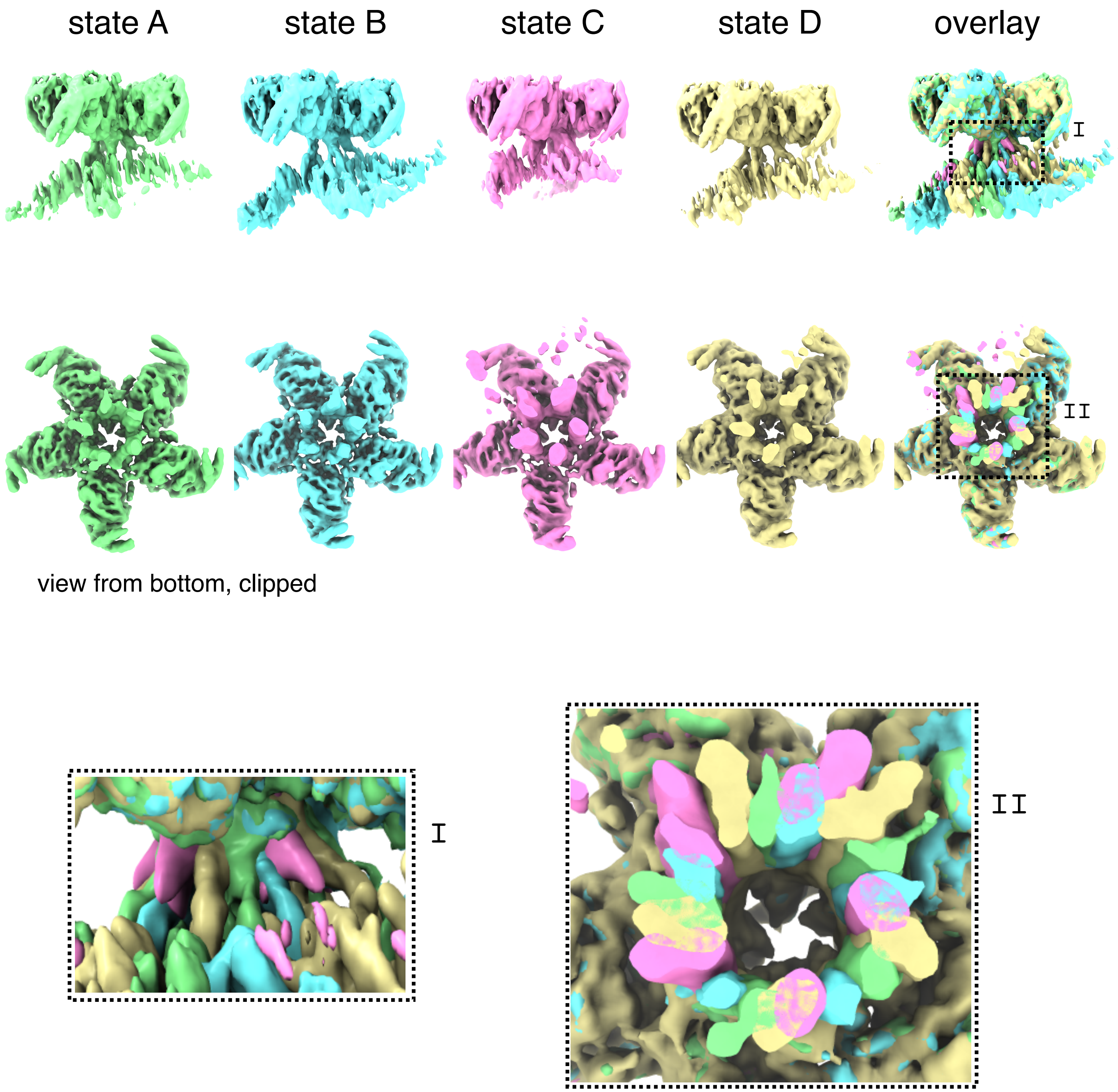


**Fig. S12.** Analysis of the cryo-em density maps in the linker region. All maps were aligned to a common setting of the "top" region and contoured at a level to emphasize the linker region. Top row: side views of maps for states A, B, C, D. Middle row: same maps rotated 90° and clipped to the middle linker region. Bottom row: zoomed in views of boxes `I` and `II` from the overlays.

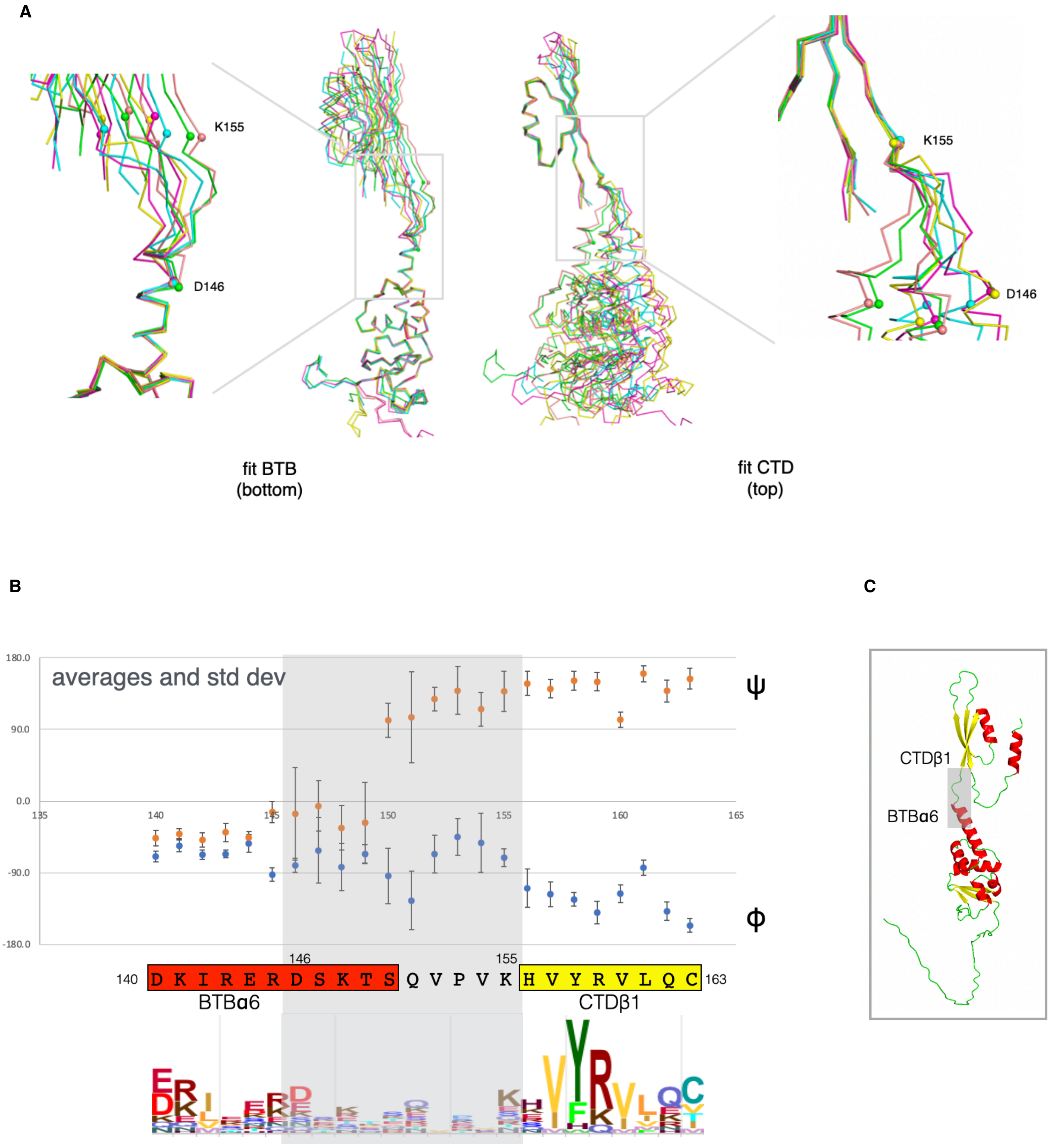


**Fig. S13.** Analysis of the hinge region between KCTD5$^{BTB}$ and KCTD5$^{CTD}$ based on two crystal structures of pentameric KCTD5 (PDB ID 3DRX and 3DRY; 10 chains in total). A. Structural alignments of the BTB domain alone (left) or the CTD alone (right) demonstrate that the conformational variations in the chains are localized to the hinge region from residues D146 to K155. B. Analysis of the variability in the backbone dihedral angles near the hinge. The average and standard deviations for the (ϕ,ψ) angles of the backbone are plotted. Standard deviations are large in the hinge region relative to the non-hinge region. A sequence logo of KCTD5 homologs showing the sequence conservation in the hinge region is shown below the human KCTD5 sequence. C. AlphaFold structure of KCTD5 with the hinge region indicated in the grey box.

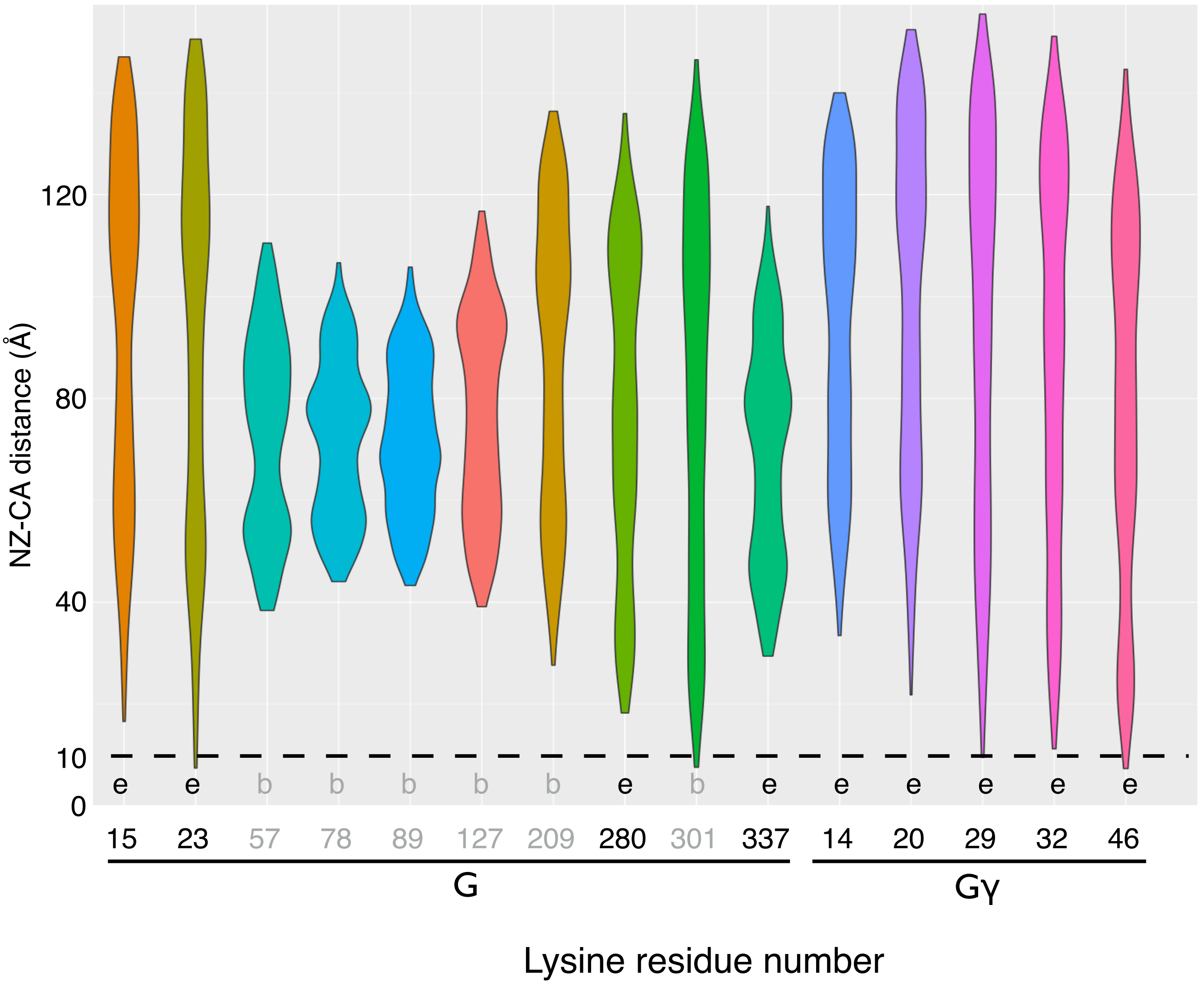


**Fig. S14.** Distance distributions between the epsilon-amino groups of the Gβ and Gγ lysine residues and the modelled Cα positions of ubiquitin G75 in the ensemble of structures. "e" and "b" indicate whether the Gβγ residues are surface-exposed or buried. These distance distributions do not take into account the flexibility in the Cullin/RBX1/ARIH1(Rcat)/Ub domain.

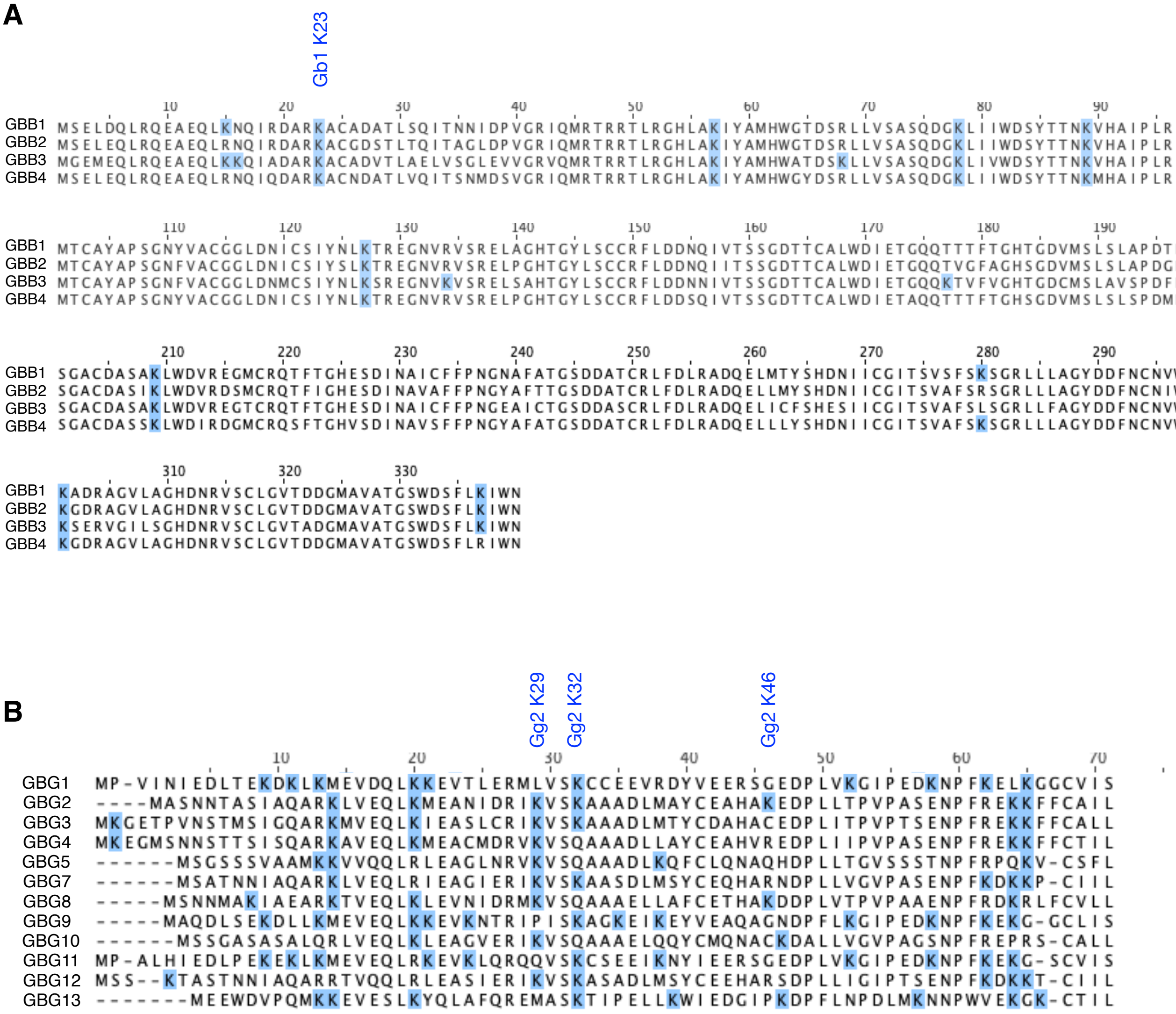

**Fig. S15.** Multiple sequence alignments of Gβ and Gγ proteins. (A) Multiple sequence alignment of human Gβ1-4. The atypical Gβ5 protein (not included) is more distantly related and binds to R7 RGS proteins instead of Gγ proteins. (B) Multiple sequence alignment of the 12 human Gγ proteins. Residue numbering for Gγ is based on Gγ2.

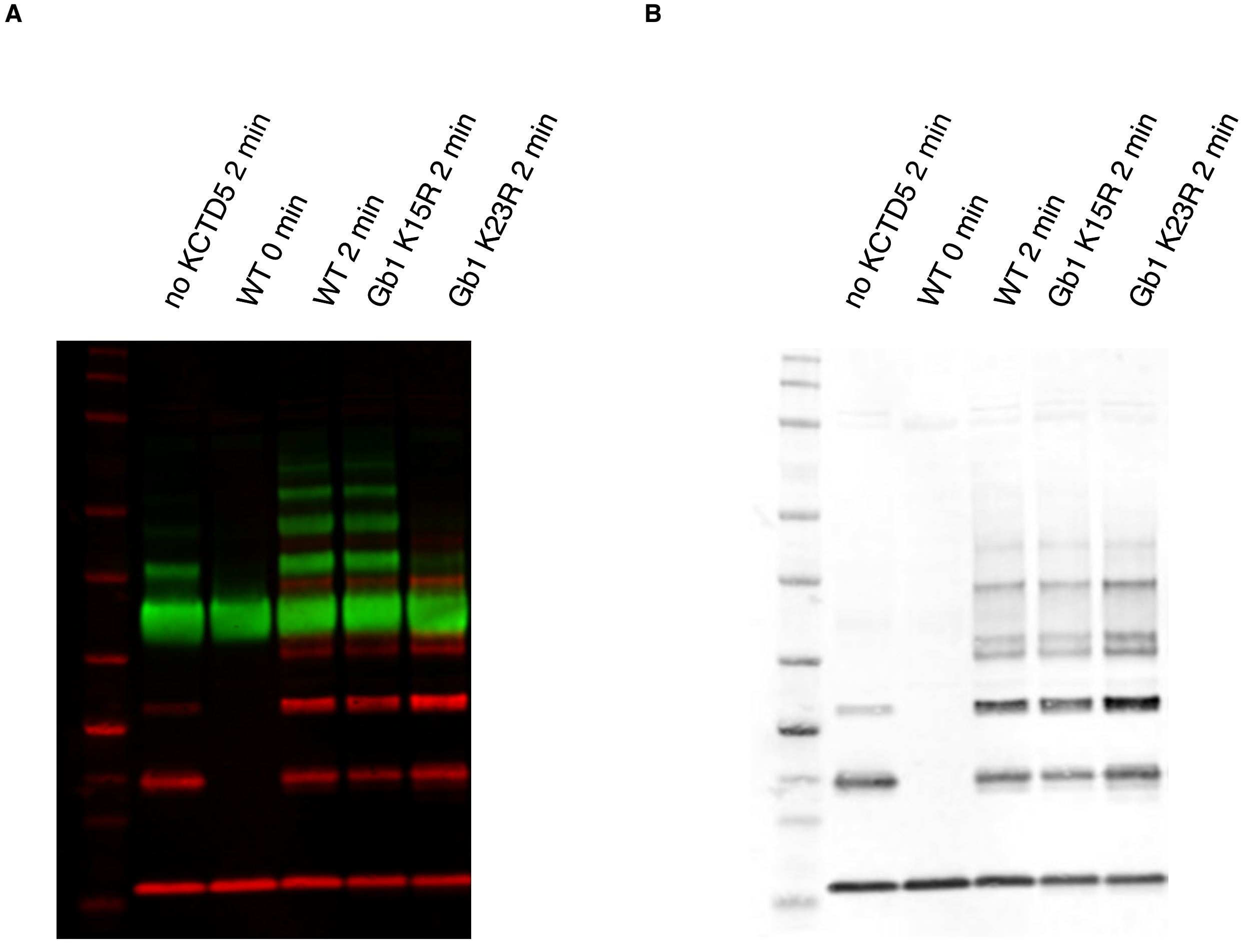


**Fig. S16.** Ubiquitylation of Gβ with mutations of candidate lysine residues. (A) Western blot image (anti-Gβ green, anti Gγ red). (B) Greyscale image of the Gγ channel. The greyscale image for the Gβ channel is shown in the main text Fig. 4.

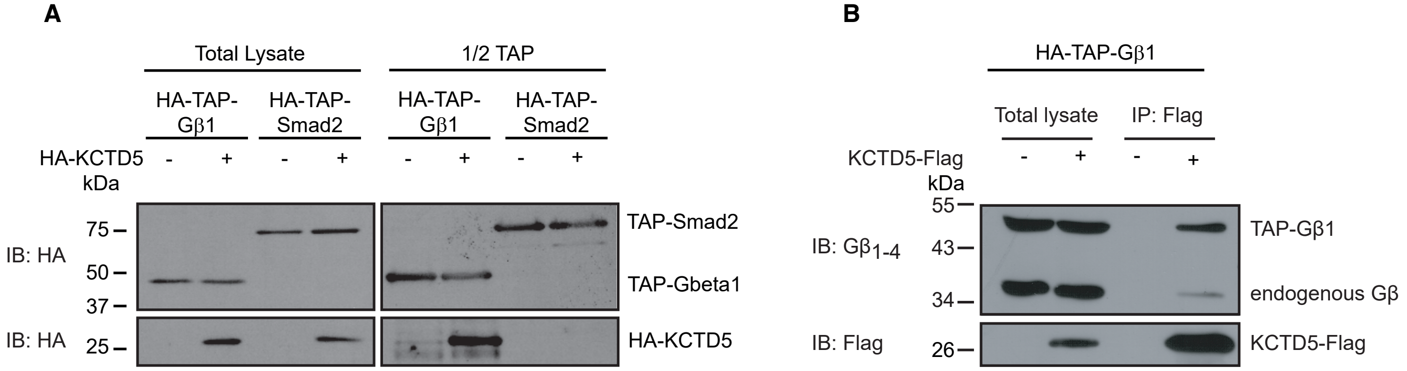


**Fig. S17.** Affinity purification and co-immunoprecipitation confirm Gβ1-KCTD5 interaction in HEK 293 cells stably expressing HA-TAP-Gβ1. (A) Lysate from pools of HA-TAP-Gβ1 or HA-TAP-Smad2 (negative control) stable cells transfected with an HA-KCTD5 DNA construct were subjected to streptavidin purification (1/2 TAP) followed by Western blot analysis. Blots are representative of 2 independent experiments. (B) Immunoprecipitation of Flag-tagged KCTD5 followed by co-detection of HA-TAP-Gβ1 and endogenous Gβ1-4. Blots are representative of 3 independent experiments.

**Table. S1.** BLI data

| target | analyte | $k_{on}$ ($10^4$ $M^{-1}s^{-1}$) | $k_{dis}$($10^{-2}$ $s^{-1}$) | $K_D$($10^{-6}$M) (kinetic) | $K_D$($10^{-6}$M) (steady state) |
|---|---|---|---|---|---|
| KCTD5 | Gβɣ | 1.9(2) | 3.7(4) | 1.98(3) | 1.80(4) |
| KCTD5/Cul3$^{NTD}$ | Gβɣ | 1.3(1) | 3.2(2) | 2.50(3) | 2.20(4) |
| KCTD5(F128R) | Gβɣ | 2.5(2) | 2.5(2) | 1.18(1) | 1.10(2) |
| KCTD5(L161R) | Gβɣ | n.d. | n.d | n.d. | n.d. |
| KCTD5(F128R/L161R) | Gβɣ | n.d. | n.d. | n.d. | n.d. |
| KCTD5(L209*) | Gβɣ | 1.4(1) | 33(3) | 23.4(3) | 13.0(1) |
| KCTD5 | Gβ(K78E)ɣ | n.d. | n.d. | n.d. | n.d. |
| KCTD5 | Cul3$^{NTD}$ | 1.9(9) | 0.05(1) | 0.03(1) | 0.03(1) |
| KCTD5(F128R) | Cul3$^{NTD}$ | n.d. | n.d. | n.d | n.d. |
| KCTD5(L161R) | Cul3$^{NTD}$ | 2.4(8) | 0.06(1) | 0.03(1) | 0.03(1) |
| KCTD5(F128R/L161R) | Cul3$^{NTD}$ | n.d. | n.d. | n.d. | n.d. |
| KCTD5(L209*) | Cul3$^{NTD}$ | 3(2) | 0.05(1) | 0.03(2) | 0.03(2) |

KCTD5 is ALFA-His-Avi-KCTD5(1-234), Gβɣ is Gβ1ɣ2(C68S), and Cul3$^{NTD}$ is Cul3(1-381).
KCTD5 mutant F128R is the Trx-His6-Avitag-KCTD5 background, while mutants L161R, F128R/L161R and L209* are in a His8-SUMO-Avitag-KCTD5 background. L209* is a mutant with a stop codon at codon 209. Values are the mean of at least three experiments, with the standard deviation indicated in the last digit. n.d.: not determined.

**Table. S2.** Cryo-em statistics

| | top only | bottom only | State A | State B | State C | State D |
|---|---|---|---|---|---|---|
| | KCTD5$^{CTD}$<br>Gβɣ | KCTD5$^{BTB}$<br>CUL3$^{NTD}$ | KCTD5<br>CUL3$^{NTD}$<br>Gβɣ | KCTD5<br>CUL3$^{NTD}$<br>Gβɣ | KCTD5<br>CUL3$^{NTD}$<br>Gβɣ | KCTD5<br>CUL3$^{NTD}$<br>Gβɣ |
| EMDB ID | EMD-41994 | EMD-41995 | EMD-41996 | EMD-42000 | EMD-42004 | EMD-42008 |
| PDB ID | 8U7Z | 8U70 | 8U81 | 8U82 | 8U83 | 8U84 |
| Data collection and processing | | | | | | |
| # movies collected (untiled/tilted/total) | 4,631 / 4,039 / 8,670 | | | | | |
| # cleaned movies (untiled/tilted/total) | 4,367 / 3,097 / 7,464 | | | | | |
| # Initial particle images (untiled/tilted/total) | 2,337,360 / 1,766,602 / 4,103,962 | | | | | |
| # cleaned particle images (untiled/tilted/total) | 135,662 / 67,200 / 202,862 | | | | | |
| # final particle images | 202,862 | 64,612 | 48,980 | 48,895 | 57,012 | 47,975 |
| Applied symmetry | C1 | C1 | C1 | C1 | C1 | C1 |
| Map resolution estimate (Å) | 2.9 | 3.5 | 3.8 | 3.8 | 4.0 | 3.9 |
| (FSC half maps; 0.143; masked) | | | | | | |
| Model | | | | | | |
| # Chains | 15 | 10 | 20 | 20 | 20 | 20 |
| # Atoms | 18,705 | 38,482 | 37,895 | 37,895 | 37,890 | 37,895 |
| # Residues | 2,420 | 2,360 | 4,770 | 4,770 | 4,770 | 4,770 |
| Bonds (RMSD) | | | | | | |
| Length (Å) | 0.005 | 0.005 | 0.005 | 0.004 | 0.019 | 0.005 |
| Angles (°) | 1.18 | 0.69 | 1.053 | 0.626 | 1.435 | 1.076 |
| MolProbity score | 2.06 | 2.00 | 2.05 | 2.16 | 2.34 | 2.04 |
| Clash score | 16.24 | 15.29 | 15.61 | 19.25 | 28.42 | 14.32 |
| Ramachandran plot (%) | | | | | | |
| Outliers | 0.04 | 0.00 | 0.13 | 0.00 | 0.36 | 0.25 |
| Allowed | 4.90 | 4.44 | 4.97 | 5.62 | 5.52 | 5.20 |
| Favored | 95.06 | 95.56 | 94.9 | 94.38 | 94.12 | 94.55 |
| Model vs. Data CC | 0.87 | 0.80 | 0.88 | 0.90 | 0.89 | 0.90 |

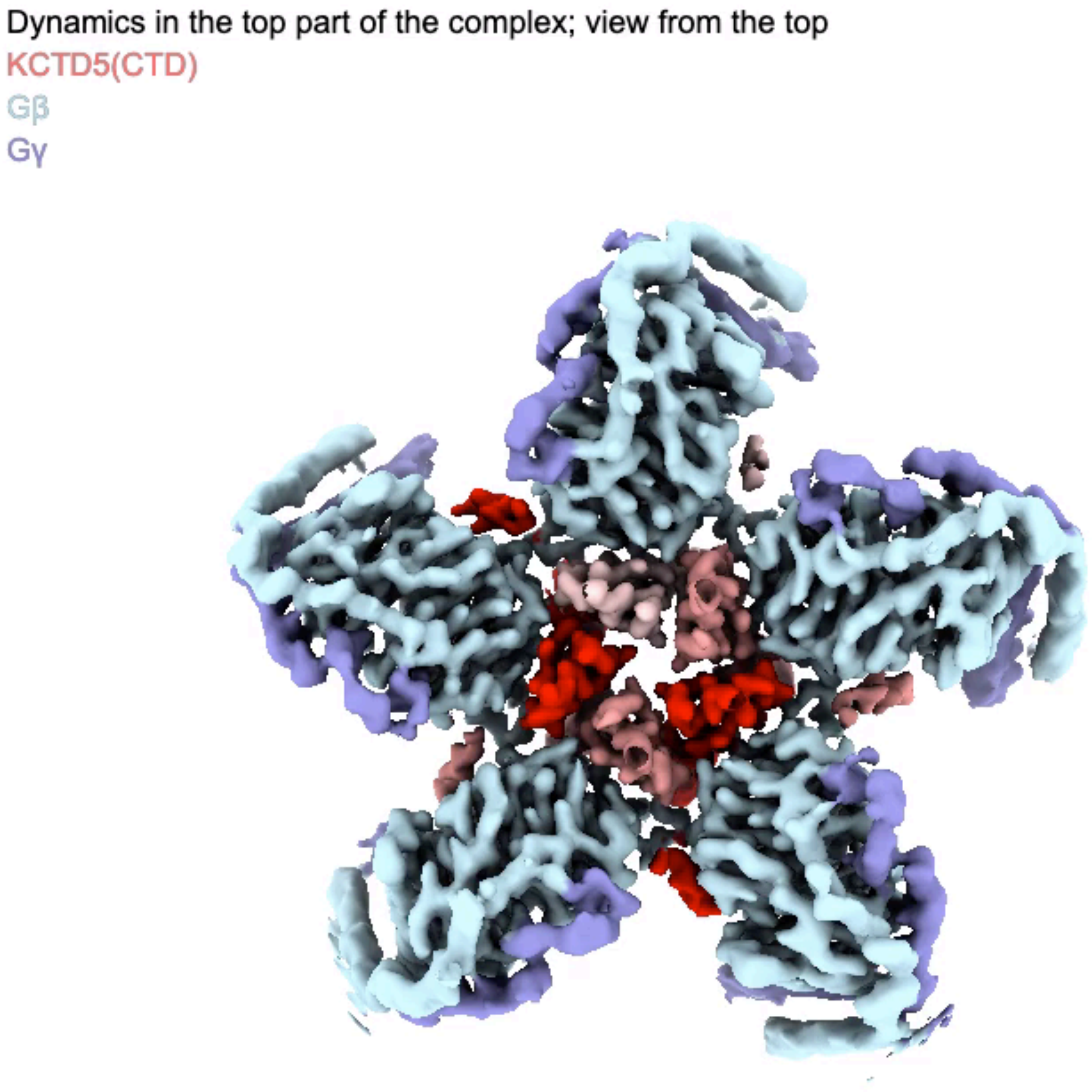


**Movie 1** Cryo-em maps of the dynamics in the top and bottom parts of the complex as revealed by 3DFlex analysis.

Dynamics in the core complex

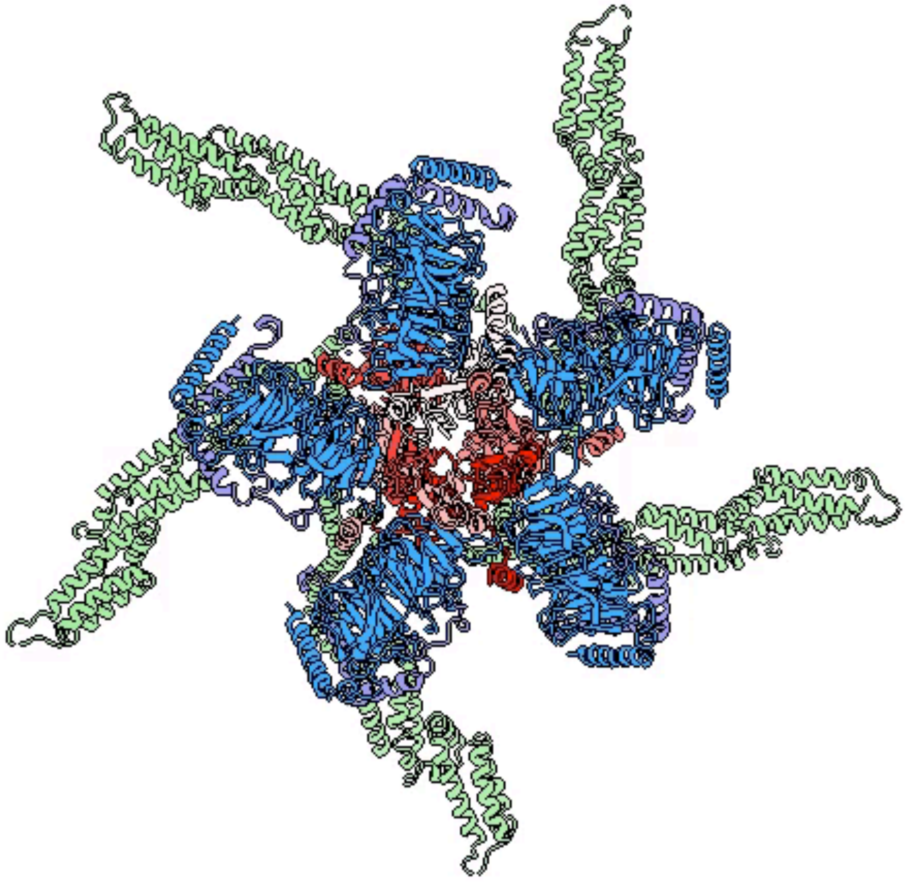

**Movie 2.** Dynamics in the CRL3$^{KCTD5}$/Gβγ complex, including the extended model that includes full-length CUL3 (green), RBX1 (grey), ARIH1 (white) and ubiquitin (yellow). The C-terminus of ubiquitin is shown as a sphere.
